# Human tonsil organoids reveal innate pathways modulating humoral and cellular responses to ChAdOx1

**DOI:** 10.1101/2024.10.11.617829

**Authors:** Maria Fransiska Pudjohartono, Kate Powell, Eleanor Barnes, Paul Klenerman, Nicholas M Provine

## Abstract

The COVID-19 pandemic response demonstrated the effectiveness of adenovirus vector vaccines in inducing protective cellular and antibody responses. However, we still lack mechanistic understanding of the factors regulating immunity induced by this platform, especially innate pathways. We utilized a human tonsil organoid model to study the regulation of adaptive responses to ChAdOx1 nCoV-19.

Innate activation and cytokine release occurred within 24 hours and T and B cell activation and antigen-specific antibody secretion occurred during the ensuing 14-day culture. Among the immune cell populations, plasmacytoid dendritic cells (pDCs) exhibited the highest ChAdOx1 transduction levels. pDC-derived IFN-α was critical for humoral responses, but production of antigen in pDCs was dispensable. Furthermore, IL-6 enhanced humoral responses in both IFN-α-dependent and independent manners, indicating intricate signaling interplay. IFN-α and IL-6 also regulated the function of vaccine-activated CD4^+^ T cells, including T_FH_. These data provide key insights into innate pathways regulating ChAdOx1-induced immunity and highlights the promise of this model for vaccine platform mechanistic studies.

**SUMMARY:** Despite the success of adenovirus vector vaccines, we still lack understanding of the mechanisms regulating immunity induced by this platform. This study utilized human tonsil organoids to provide key insights into innate pathways regulating ChAdOx1-induced humoral and cellular immunity.

## INTRODUCTION

The COVID-19 pandemic response expedited the implementation of new vaccine platforms, including adenovirus vector vaccines. Several adenovirus vector vaccines were introduced against SARS-CoV-2, with effectiveness in inducing antibodies to SARS-CoV-2 and protecting from COVID-19 disease (Voysey *et al*., 2021; Halperin *et al*., 2022; Sadoff *et al*., 2022). However, despite the successful deployment of this vaccine, mechanistic understanding of the factors governing the humoral responses to adenovirus vector vaccines is still lacking, with most studies focused on using the technology to induce cellular immunity (Provine and Klenerman, 2022). Thus, there is a major gap in our understanding of this vaccine platform.

Beyond a few select studies, little is known about the regulation of humoral immunity to the transgene antigen cargo of adenoviral vector vaccines (Provine and Klenerman, 2022). Protection against COVID-19 is associated with the production of IgG antibodies targeting SARS-CoV-2, which requires the development of antigen-specific plasma cells and memory B cells from germinal centers in lymph nodes. In the germinal center (GC) reaction, CD4^+^ T cells and particularly follicular helper T cells (T_FH_) play an important role in providing costimulatory signaling to B cells. The induction of antibodies towards the transgene of Ad vaccines is known to be primarily driven by GCs (Foster *et al*., 2022), requiring CD4 T cells (Provine *et al*., 2016) and specifically T_FH_ (Foster *et al*., 2022).

However, less is known about the innate signals that initiate the GC responses to adenoviral vaccines. Innate immune responses can modulate the GC response and outcomes, with innate immune phenotypes being associated with adaptive immune response outcomes to other vaccine platforms (Fourati *et al*., 2022; Hagan *et al*., 2022). For adenoviral vaccines, TLR4 signaling has been shown to promote antibody responses (Li *et al*., 2018). One study has found that induction of antibodies against the adenovirus viral particle itself is dependent on type I interferon, which regulated multiple steps of B cell differentiation (Zhu, Huang and Yang, 2007). However, it is unclear if similar signals regulate the immune response to the transgene antigen. A recent study in NHPs found a correlation between type I IFN levels and antibody titers against a SARS-CoV-2 spike transgene (He *et al*., 2022), suggesting an association. Direct experimental validation is required, and more generally, a broad survey of critical innate immune pathways is needed to unravel the biology of these vaccines.

To investigate the role of these innate-adaptive immune cell interactions in modulating humoral responses, it is critical to model these processes occurring at the site of vaccine responses – secondary lymphoid tissues. A recently described in vitro human lymphoid tissue organoid model utilizing tonsil tissue has shown utility for investigating the regulation of human B cell responses (Wagar *et al*., 2021; Kastenschmidt *et al*., 2023; Yin *et al*., 2023; Mitul *et al*., 2024). Providing a model for studying both antibody and cellular biology in an antigen-specific manner, the system has been used quite heavily to study vaccine responses to influenza A virus vaccines. The manipulability of the model has allowed description of the role for specific cell types in live-attenuated influenza virus vaccine responses (Wagar *et al*., 2021), and differential roles for memory or naive B cell populations based on live-attenuated or inactivated vaccine formulation (Kastenschmidt *et al*., 2023). It has been demonstrated that adenoviral vector induced antibody responses can be measured, but no specific investigation of the regulation of these responses was performed (Wagar *et al*., 2021). Thus, this organoid model has promise for unravelling pathways regulating adenoviral vector induced humoral responses.

We utilized this human in vitro tonsil organoid model to investigate the processes that regulate the induction of T cell, B cell and antibody responses to the ChAdOx1 nCoV-19 vaccine. The tonsil organoid system demonstrated cellular, humoral and innate responses to adenovirus vaccines, presenting a viable model for mechanistic studies. We discovered that plasmacytoid dendritic cells modulate the humoral response to adenovirus vector vaccines, mainly through secretion of type 1 IFN. Type 1 IFN augments antibody production through modulating both innate and adaptive responses, both directly and indirectly through IL-6 induction. These two cytokines also worked in concert to regulate the phenotype and function of vaccine-activated CD4^+^ T cells.

## RESULTS

### Adenovirus vector vaccines induce humoral responses in tonsil organoid cultures

Our first aim was to validate that the tonsil organoid culture system could robustly model humoral responses to adenovirus vector vaccines. To achieve this, we stimulated tonsil organoid cultures with ChAdOx1 nCoV-19 over 14 days of culture (experimental schematic in Figure S1A). Viability and cell recovery remained consistent over the culture period (Figure S1B).

To validate that the system could induce specific antibody production in response to vaccine stimulation, anti-SARS-CoV-2 S1 IgG was measured in the culture supernatants. Anti-S1 IgG production was very low (<10 ng/mL) or undetectable in unstimulated cultures from 18/22 post-pandemic donors (Figure 1A). Peculiarly, in the post-pandemic donors that produced anti-S1 IgG when unstimulated, ChAdOx1 stimulation suppressed specific antibody production (Figure S1C). Given their unsuitability for addressing the research question, these samples were excluded from subsequent analyses. ChAdOx1 nCoV-19 stimulation induced robust S1-specific antibody responses in the majority of post-pandemic donors (n=13/16; Figure 1A). Stimulation of tonsils collected prior to the COVID-19 pandemic (“pre-pandemic”) did not induce S1-specific IgG production (Figure 1B). To test for non-specific IgG induction by the ChAdOx1 vector, ChAdOx1 GFP was used as a control. ChAdOx1 GFP did not induce anti-S1 IgG production in any of the 4 post- and 2 pre-pandemic donors tested (Figure 1A).

**Figure 1:**
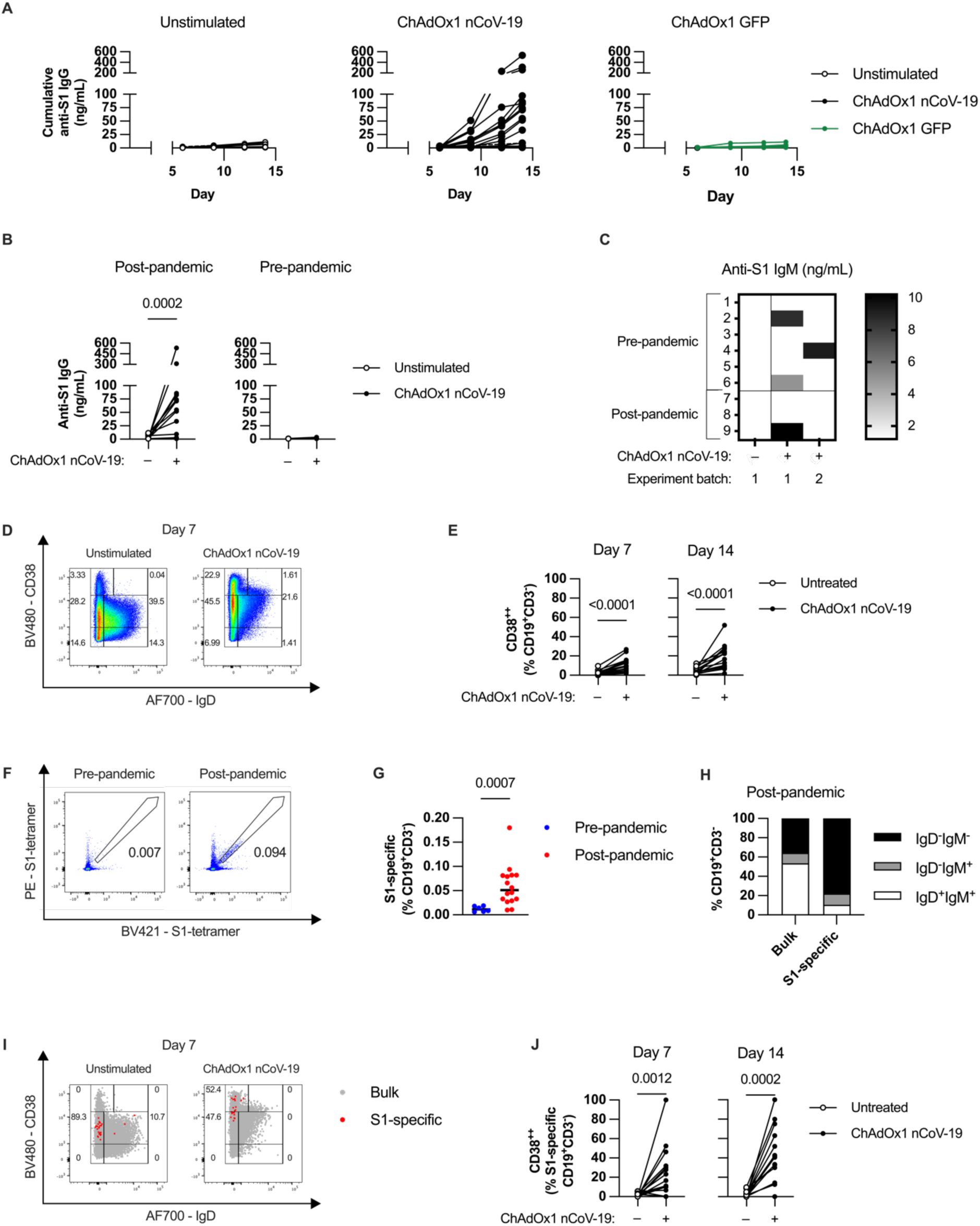
ChAdOx1 nCoV-19 induces specific antibody production and B cell activation in post-pandemic tonsil organoids. **(A)** Cumulative anti-S1 IgG production over time in unstimulated tonsil organoids (*left*) or in response to ChAdOx1 nCoV-19 (*middle*) or ChAdOx1 GFP (*right*). **(B)** Cumulative anti-S1 IgG production with ChAdOx1 nCoV-19 stimulation in tonsils collected post- and pre-COVID-19 pandemic. **(C)** Cumulative anti-S1 IgM production on day 14 in pre- and post-pandemic tonsil organoids over repeated experiments. **(D)** Representative FACS plot of B cells (singlet, live, CD45^+^CD19^+^CD3^-^) from unstimulated and ChAdOx1-stimulated organoids on day 7 of culture. **(E)** Bulk B cell activation measured by % of CD38^++^ cells in response to ChAdOx1 nCoV-19 in post-pandemic donors. **(F)** Representative FACS plot of S1-tetramer binding of B cells from pre- and post-pandemic donors. **(G)** Percentage of S1-specific B cells in pre- and post-pandemic donors. **(H)** Surface IgD and IgM expression on S1-specific B cells in post-pandemic donors based. **(I)** Representative FACS plot of S1-specific B cells overlaid on bulk B cells on Day 7 of organoid culture from a post-pandemic donor. **(J)** S1-specific B cell activation measured by %CD38^++^ cells in response to ChAdOx1 nCoV-19 in post-pandemic donors. Data in A-B are combined from 5 experiments with 24 donors, C from 2 experiments with 9 donors, data in E-J from 4 experiments with 16 donors. Each symbol represents an individual donor. Values in B, E and J were compared using Wilcoxon matched pairs signed rank test, values in G were compared using Mann-Whitney test.

We hypothesized that the lack of S1-specific antibodies in pre-pandemic donors might reflect insufficient time in the culture for class-switching to occur. Thus, we examined IgM production as an earlier product of naive B cell activation. In contrast to IgG production, anti-S1 IgM was produced in both post- and pre-pandemic donors in a variable manner across two experiment batches (Figure 1C). The high variability between technical replicates suggests that specific IgM production in the model is stochastic, possibly depending on the presence in the culture well of S1-specific naïve B cells, which are present at low frequencies (Feldman *et al*., 2021).

After validating antibody production upon stimulation, we measured B cell activation using flow cytometry. We compared B cell subsets based on IgD and CD38 expression as naïve and activation markers, respectively (Sanz *et al*., 2019). ChAdOx1 nCoV-19 decreased the percentage of IgD^+^ B cells and increased CD38 expression on B cells (Figure 1D). Overall, the most striking change was an increase in IgD^-^CD38^++^ B cells, a population commonly identified as antibody secreting cells (ASCs) (Sanz *et al*., 2019), which encompasses both plasmablasts and plasma cells (Figure 1D-E). This population had higher CD27 expression compared to other subsets (Figure S1D), in line with previous observations for ASCs (Sanz *et al*., 2019). A similar increase in IgD^-^CD38^++^ B cells was observed for ChAdOx1 GFP (Figure S1E), but did not induce S1-specific antibody production (Figure 1A).

Next, we identified antigen-specific B cells using fluorescently-labelled S1 tetramers. Post-pandemic donors showed a clear population of S1-specific B cells, while pre-pandemic donors did not (Figure 1F-G). In post-pandemic donors, the S1-specific B cells were mainly composed of switched memory (IgD^-^IgM^-^) B cells compared to the bulk B cell population (Figure 1H). The percentage of S1-specific B cells decreased with ChAdOx1 nCoV-19 stimulation (Figure S1F), possibly due to activated cells downregulating their B cell receptors (BCR) as they differentiate into ASCs. However, S1-specific cells were detectable, and were highly activated in response to stimulation (Figure 1I-J). These findings suggest that S1-specific IgG production, but not IgM production, in response to stimulation is associated with detectable S1-specific memory B cells. Most importantly, the combination of these data gave us high confidence in the utility of the model for probing pathways regulating ChAdOx1-induced humoral immunity.

### Plasmacytoid dendritic cells (pDCs) contribute to the early response to ChAdOx1 nCoV-19 through antigen expression and cytokine secretion

Next, we investigated the early innate response to ChAdOx1 nCoV-19 in the tonsil organoid model after 24 hours in culture. As S1-specific IgG production was only consistently induced in organoids from post-pandemic donors, only these donors were used for mechanistic studies. We focused on identifying the cells that are transduced and activated by ChAdOx1. To identify cells transduced by the vector, we measured cells expressing either the S1 spike peptide (for ChAdOx1 nCoV-19) or GFP (ChAdOx1 GFP) by flow cytometry. No significant changes in the tonsil cell compositions were observed with freeze-thawing (Figure S2A). Following exposure, all cells showed an increase in transgene-expressing cells, with varying frequencies (Figure 2A, S2B). The highest transduction rates were seen for dendritic cells (DCs), especially plasmacytoid dendritic cells (pDCs) (Figure 2A). The number of pDCs decreased with ChAdOx1 nCoV-19 stimulation, likely due to death following transduction (Figure S2C). Both ChAdOx1 nCoV-19 and ChAdOx1 GFP demonstrated similar patterns of transduction, with higher detection rates for GFP, likely due to ease of detecting GFP expression. Monocytes were highly transduced (66-100%) but were not included in subsequent analyses as their scarcity in tonsil made quantification unreliable (0-5 cells/well). Conventional DCs (cDCs) and CD45^-^ cells (presumably fibroblasts) were also transduced with some efficiency (mean % positive for each group). This pattern of transduction is highly consistent with what has been reported in peripheral blood mononuclear cells (PBMCs) (Provine *et al*., 2021).

**Figure 2:**
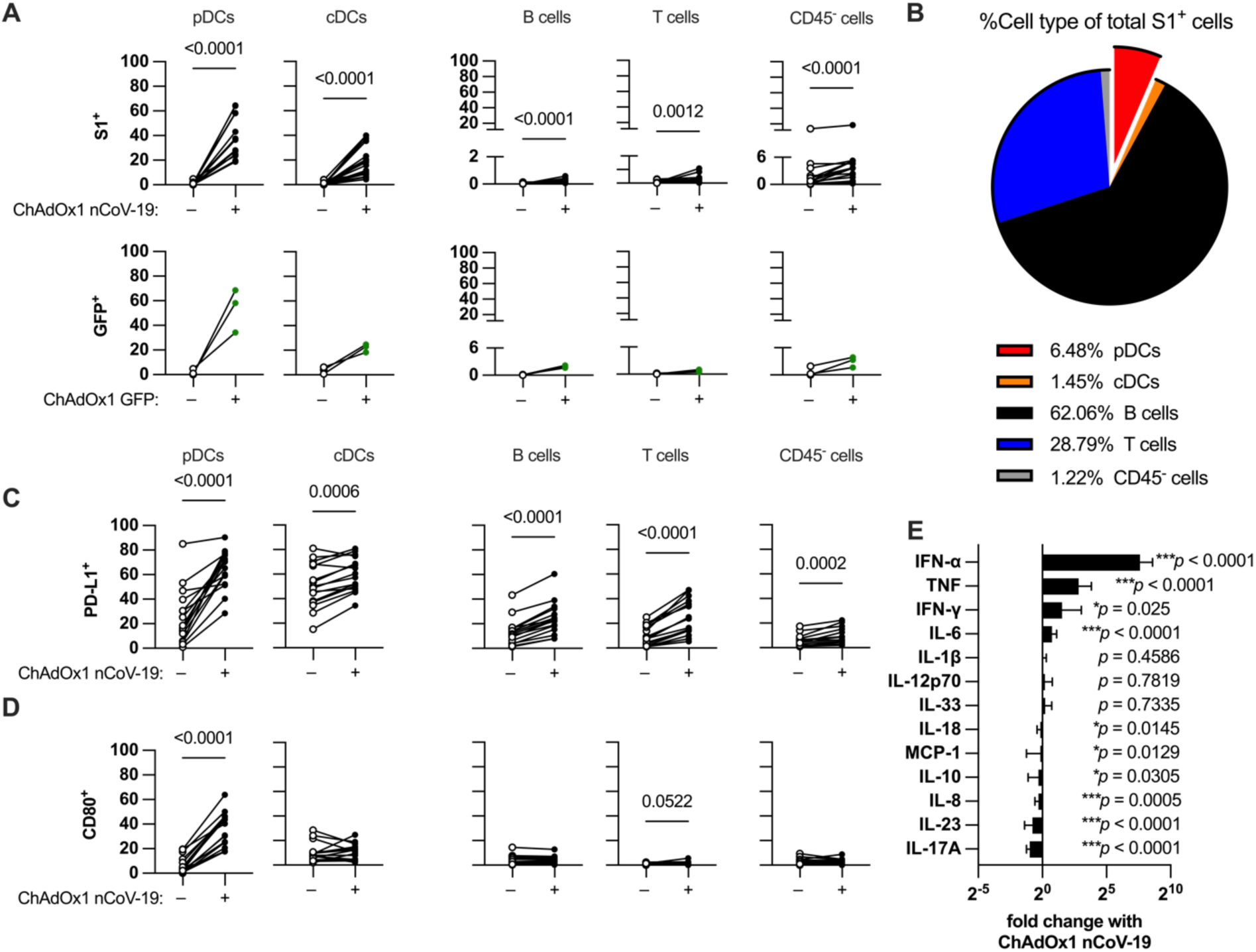
Characterization of innate immune activation and cellular transduction by ChAdOx1 in tonsil organoids. **(A)** Transduction rates of different cell types in the tonsil organoid culture system. Transduction was measured based on detection of ChAdOx1 transgene (either S1 spike or GFP) by flow cytometry. **(B)** Average fraction of each cell type as a proportion of all transduced cells. **(C-D)** Innate immune cell activation by ChAdOx1 nCoV-19 as measured by percentage of cells expressing surface PD-L1 **(C)** or CD80 **(D)**. **(E)** Relative change in cytokine levels in tonsil organoid media supernatants with ChAdOx1 nCoV-19 stimulation compared to unstimulated (each bar shows mean ± SD). All results were from cell/media harvest at 24 hours after plating. Data in A-E are combined from 4 experiments with a total of 16 donors. Each symbol represents an individual donor. Values in A and C-D were compared using Wilcoxon matched pairs signed rank test. Values in E were compared using Wilcoxon matched pairs signed rank test from unstimulated compared to ChAdOx1 nCoV-19-stimulated wells (data in Fig. S2D).

Despite having the highest transduction efficiency, pDCs only contributed a small fraction of total transduced cells (6.5%), due to the relatively small size of the pDC population in the culture (∼1% of all cells) (Figure 2B). By contrast, B cells were the main contributor of transduced cells despite very low transduction rates due to the high frequency of B cells in the culture system (∼65% of all cells).

We next evaluated immune activation through the expression of PD-L1 and CD80. ChAdOx1 nCoV-19 stimulation upregulated PD-L1 expression on all cell populations, while CD80 was only induced on pDCs (Figure 2C-D). Other than pDCs, most immune cell types had much higher PD-L1 expression rates than their transduction rates, suggesting indirect activation of non-transduced cells, likely through cytokines.

We measured ChAdOx1 nCoV-19 induced cytokine secretion at 24 hours. ChAdOx1 nCoV-19 induced production of several pro-inflammatory cytokines, including IFN-α, TNF, IFN-γ, and IL-6, with the greatest induction of IFN-α (Figure 2E, Figure S2D). The induced cytokines are consistent with previous studies on patient samples and PBMC stimulation using ChAdOx1 nCoV-19 (Jiang *et al*., 2023).

### pDCs modulate the humoral response to ChAdOx1 nCoV-19, primarily through type 1 IFN

Given the high transduction and activation of pDCs as well as the IFN-α induced, we hypothesized that pDCs were an important early immune modulator of response to adenovirus vector vaccines. Thus, we performed the organoid culture with pDC depletion (see methods). The depletion process removed 98.6-99.9% of pDCs, while the percentage of other cells remained stable except for a slight increase of CD45^-^ cells (Figure S3A).

pDC-depletion dramatically impaired S1-specific IgG induction in all responsive donors, confirming the modulatory function of pDCs on antibody production (Figure 3A). In accordance with the decreased antibody production, both bulk and S1-specific B cell activation was also dampened by pDC-depletion (Figure 3B-C). pDC-depletion did not significantly affect the overall antigen availability as shown by the total frequency of transduced cells in the culture (Figure 3D) nor did it alter transduction rates for other cell types (Figure 3E). However, early activation of other immune cells (cDCs, B cells, T cells), as assessed by PD-L1 expression, decreased with pDC-depletion (Figure 3F). There was minimal impact on CD80 expression, as no cells showed clear induction of CD80 (Figure S3B). As expected, pDC-depletion dramatically reduced levels of IFN-α (Figure 3G). However, it also resulted in a reduction of the other induced pro-inflammatory cytokines TNF, IFN-γ, and IL-6. These results demonstrate that pDCs are critical modulators of the humoral responses to adenovirus vectors.

**Figure 3:**
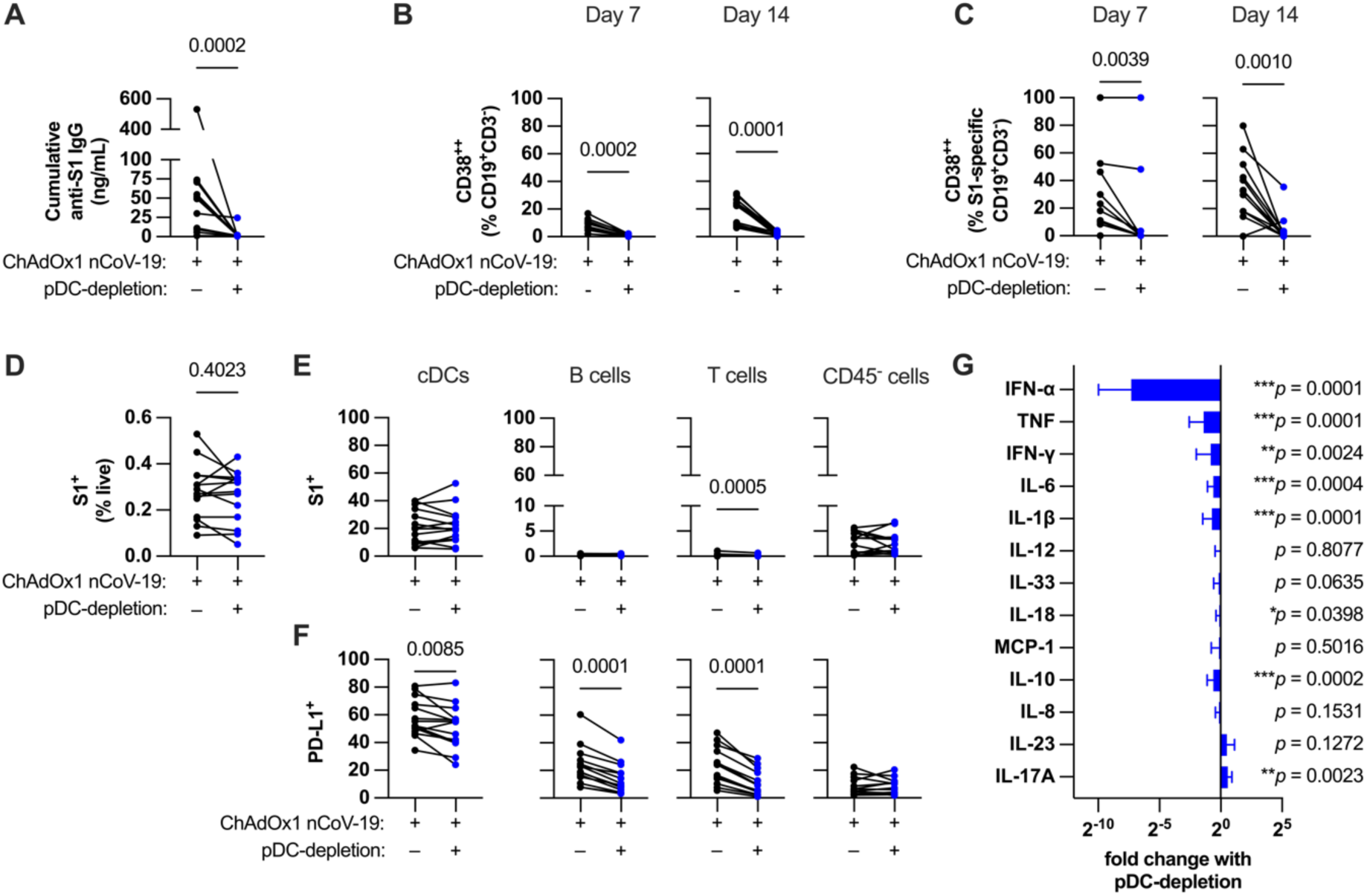
pDC-depletion impairs both humoral and innate responses to ChAdOx1 nCoV-19. **(A)** Cumulative anti-S1 IgG production in ChAdOx1 nCoV-19-stimulated post-pandemic tonsil organoids with or without pDC-depletion after 14-day culture. **(B, C)** Bulk **(B)** and S1-specific **(C)** B cell activation in ChAdOx1 nCoV-19-stimulated tonsil organoids with or without pDC-depletion. **(D)** Percentage of total transduced cells after 24-hour culture in ChAdOx1 nCoV-19-stimulated tonsil organoids with or without pDC-depletion. **(E)** Transduction rates of different cell types in ChAdOx1 nCoV-19-stimulated tonsil organoids with or without pDC-depletion. **(F)** Innate activation of different cell types based on surface PD-L1 in ChAdOx1 nCoV-19-stimulated tonsil organoids with or without pDC-depletion. **(G)** Relative change in cytokine levels in ChAdOx1 nCoV-19-stimulated tonsil organoid supernatants with or without pDC-depletion (each bar shows mean ± SD). Data in A-G are combined from 4 experiments with a total of 14 donors. Each symbol represents an individual donor. Values in A-F were compared using Wilcoxon matched pairs signed rank test. Values in G were compared using Wilcoxon matched pairs signed rank test from ChAdOx1 nCoV-19-stimulated untreated wells compared to stimulated wells with pDC-depletion (data in Fig. S3C).

### IFN-α is the main mediator of pDC-mediated modulation of the humoral response

To investigate whether the effect of pDCs on the humoral response was mediated by type 1 IFN, as a source of antigen, or both, we investigated the effect of blocking type 1 IFN on the culture. Type 1 IFN blockade was sufficient to phenocopy pDC-depletion, significantly reducing S1-specific IgG secretion (Figure 4A) and bulk B cell activation (Figure 4B) and S1-specific B cell activation (Figure 4C) in all donors.

**Figure 4:**
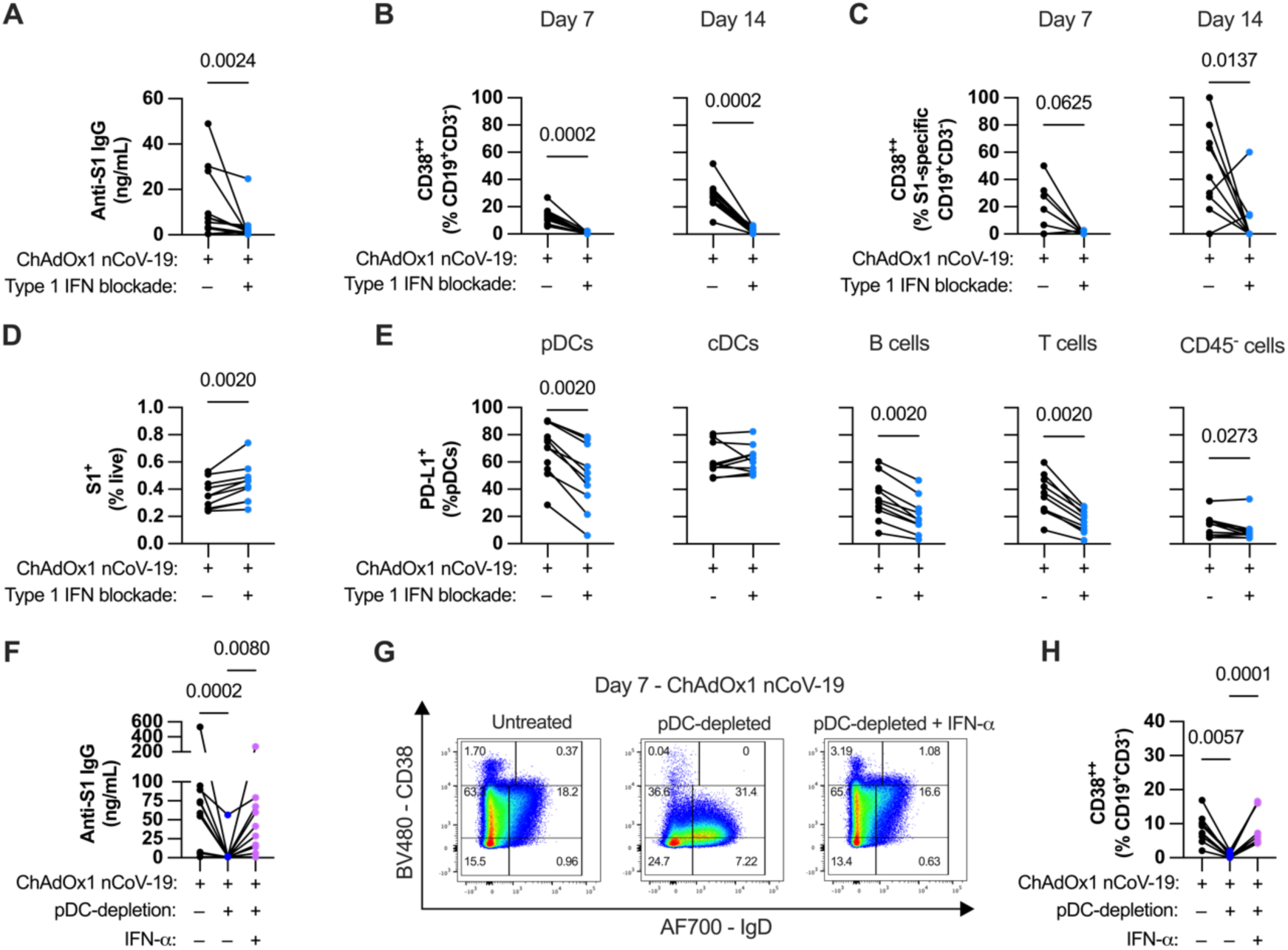
pDCs modulate the humoral response to adenoviral vectors primarily through type 1 IFN. **(A)** Cumulative anti-S1 IgG production in ChAdOx1 nCoV-19-stimulated tonsil organoids with or without type 1 IFN blockade after 14-day culture. **(B & C)** Bulk **(B)** and S1-specific **(C)** B cell activation levels in ChAdOx1 nCoV-19-stimulated tonsil organoids with or without type 1 IFN blockade. **(D)** Percentage of total transduced cells after 24-hour culture in ChAdOx1 nCoV-19-stimulated tonsil organoids with/without type 1 IFN blockade. **(E)** Innate activation of different cell types based on surface PD-L1 in ChAdOx1 nCoV-19-stimulated tonsils organoids with or without type 1 IFN blockade. **(F)** Cumulative anti-S1 IgG production in ChAdOx1 nCoV-19-stimulated tonsil organoids with pDC-depletion and with or without IFN-α supplementation. **(G)** Representative FACS plot of bulk B cells in pDC-depleted ChAdOx1 nCoV-19-stimulated tonsil organoid cultures and with or without IFN-α supplementation. **(H)** Bulk B cell activation levels in pDC-depleted ChAdOx1 nCoV-19-stimulated tonsil organoids and with or without IFN-α supplementation. Data in A-C are combined from 3 experiments with a total of 13 donors, D-E from 3 experiments with 10 donors, F and H from 3 experiments with 11 donors. Each symbol represents an individual donor. Values in A-E were compared using Wilcoxon matched pairs signed rank test, values in F and H using Friedman test with Dunn’s multiple comparisons test.

With regards to innate responses, blocking type 1 IFN increased the overall number of transduced cells (Figure 4D). The increase appears to be mediated by a particular increase in pDC transduction rates and a modest increase for B cells (Figure S4A). Interestingly, despite increased transduction rates, PD-L1 expression on most cell types decreased with type 1 IFN blockade (Figure 4E). This suggests that upregulation of surface PD-L1 expression is mediated by autocrine type 1 IFN and that transduction alone is insufficient to activate the pDCs. Induction of TNF, IFN-γ and IL-6 were marginally but consistently decreased with type 1 IFN blockade (Figure S4B), again similar to pDC depletion.

While these data clearly supported a model where pDC-derived IFN-α is important for ChAdOx1 nCoV-19-induced humoral responses, it was unclear if pDC-derived antigen was also important. To investigated this, we determined if addition of exogenous IFN-α was sufficient to rescue pDC-depleted cultures. Adding exogenous recombinant IFN-α to pDC-depleted cultures restored S1-specific IgG antibody secretion (Figure 4F) and B cell activation (Figure 4G-H). Adding IFN-α without ChAdOx1 nCoV-19 stimulation caused B cell activation (CD38^++^) but no anti-S1 IgG secretion (Figure S4C), further confirming that the cognate antigen is required for specific antibody secretion. IFN-α supplementation did not affect overall percentage of transduced cells (Figure S4D).

While cognate antigen is critical for ChAdOx1-stimulation of humoral responses (Figure 1A), these results suggest that pDCs do not modulate these responses by producing cognate antigen. Instead, they promote humoral responses primarily by producing IFN-α.

### pDC-derived type 1 IFN augments the humoral response to ChAdOx1 nCoV-19 partially by stimulating IL-6 production

Depletion of pDCs reduced IL-6 levels in the culture (Figure 2G), and type 1 IFN supplementation of pDC-depleted cultures also restored the induction of IL-6 (Figure 5A). IL-6 is a known modulator of B cell responses (Vazquez, Catalan-Dibene and Zlotnik, 2015). Thus, we sought to investigate if pDCs modulated ChAdOx1-induced humoral immunity through IL-6 as an intermediary signal. First, we tested whether IL-6 regulated humoral responses to ChAdOx1. Blockade of IL-6 signaling using anti-IL-6R antibodies significantly, but partially, decreased S1-specific IgG production across all donors (Figure 5B). There was a trend of decreased B cell activation as well, although this did not reach statistical significance (Figure 5C-D).

**Figure 5:**
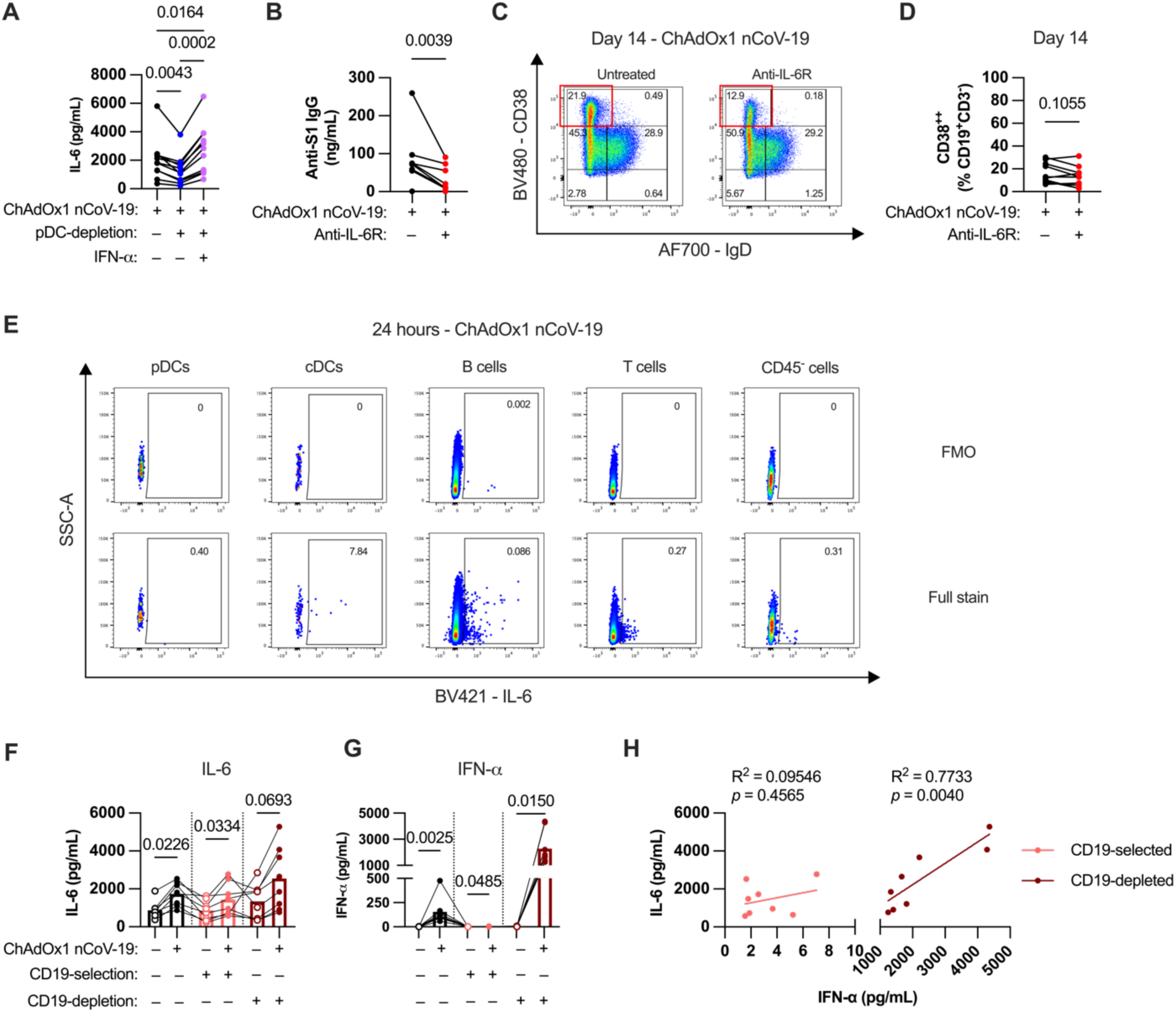
pDC-derived type 1 IFN augments the humoral response to ChAdOx1 nCoV-19 partially through driving IL-6 production. **(A)** IL-6 levels in ChAdOx1 nCoV-19-stimulated tonsil organoid supernatants with or without pDC-depletion and IFN-α supplementation. **(B)** Cumulative anti-S1 IgG production in ChAdOx1 nCoV-19-stimulated tonsil organoids with or without IL-6R blockade after 14-day culture. **(C-D)** Representative FACS plot **(C)** and group summary **(D)** of bulk B cells in tonsil organoid cultures with or without IL-6R blockade on day 14 of culture. **(E)** Representative FACS plots of IL-6 staining or FMO control of different cell types from stimulated tonsil organoids. **(F, G)** IL-6 **(F)** and **(G)** IFN-α levels in ChAdOx1 nCoV-19-stimulated tonsil organoid cultures with or without CD19-selection or CD19-depletion after 24 hours in culture. **(H)** Correlation between IL-6 and IFN-α levels in CD19-selected or CD19-depleted cultures after 24 hours in cultures with ChAdOx1 nCoV-19. Data in A are combined from 3 experiments with a total of 11 donors, B and D from 2 experiments with 9 donors, F-H from 2 experiments with 8 donors. Each symbol represents an individual donor. Values in A were compared using Friedman test with Dunn’s multiple comparisons test, values in B and D using Wilcoxon matched pairs signed rank test. Values in F-G were compared using Friedman test with Dunn’s multiple comparisons test, correlations in H were done using Spearman correlation.

As monocytes are one of the best described sources of IL-6 (Aarden *et al*., 1987), but are essentially absent in tonsils (Figure S2A), we next sought to determine the cellular source of IL-6. Thus, we performed intracellular staining of IL-6 at 24 hours after ChAdOx1 nCoV-19 stimulation. With the exception of pDCs, most cell types produced IL-6 (Figure 5E). Although the percentage of IL-6-producing B cells is low as a proportion of total B cells, given the high frequency of B cells in the culture, B cells comprise most of the IL-6-producing cells. Consistent with the flow cytometry analysis, isolated B cells produced IL-6 in response to ChAdOx1 nCoV-19 stimulation (Figure 5F). However, B cell-depleted cultures also produced substantial IL-6, suggesting that non-B cell populations are also a source (Figure 5F). Interestingly, despite both cultures producing IL-6, there were major differences in the level of IFN-α in the cultures (Figure 5G). B cell-depleted cultures had high levels of IFN-α production (consistent with the presence of pDCs) and there was a strong correlation between IFN-α levels and IL-6 levels (Figure 5H). By contrast, despite producing substantial IL-6, the purified B cell culture had negligible induction of IFN-α (<10 pg/mL) (Figure 5G), and thus IL-6 and IFN-α levels were unsurprisingly not related (Figure 5H). These findings suggest that both B cells and innate cells contribute to IL-6 induction by ChAdOx1 nCoV-19, but through type 1 IFN-independent and -dependent mechanisms, respectively.

### Type 1 IFN and IL-6 augment the cellular response to ChAdOx1 nCoV-19, including for follicular helper T (T_FH_) cells

Another important factor that could modulate humoral responses is the CD4^+^ T cell response. Thus, we sought to examine T cell activation in response to the vaccine. In the tonsil organoid model, ChAdOx1 nCoV-19 induced CD4^+^ T cell activation (CD40L^+^CD69^+^) (Figure 6A-B). ChAdOx1 nCoV-19 induced production of IFN-γ, TNF, and IL-2, as measured by intracellular cytokine staining (Figure 6C, Figure S5A). Thus, we could assess the activation and effector function of spike-specific T cells in response to ChAdOx1 nCoV-19.

**Figure 6:**
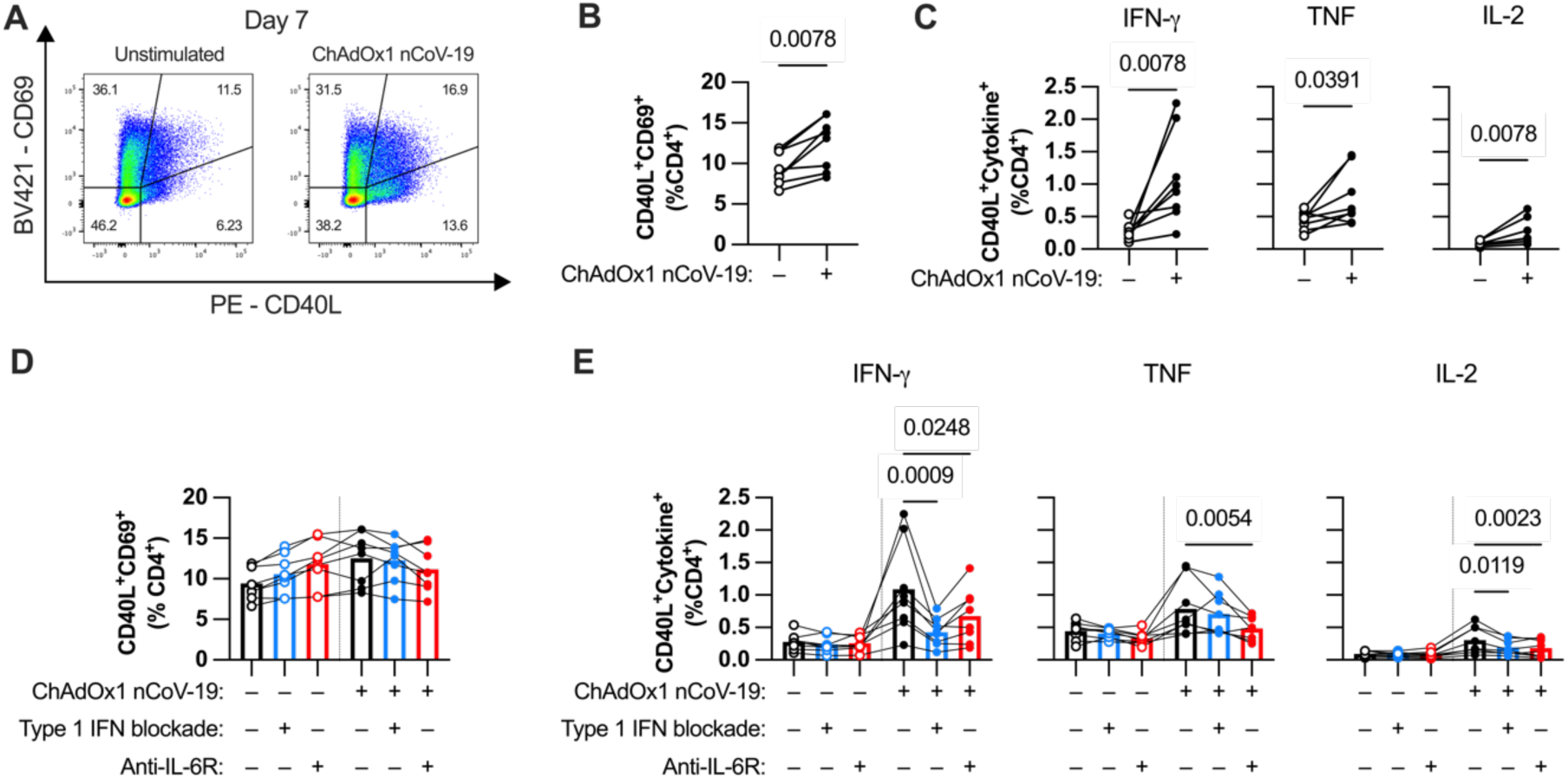
Type 1 IFN and IL-6 augment the CD4^+^ T cell response to ChAdOx1 nCoV-19. **(A-B)** Representative FACS plots of CD69 and CD40L expression on CD4^+^ T cells and group summary **(B)** in ChAdOx1 nCoV-19-stimulated tonsil organoids on day 7 of culture. **(C)** Percentage of CD4^+^ T cells co-expressing CD40L and cytokines (IFN-γ, TNF, and IL-2) in ChAdOx1 nCoV-19-stimulated tonsil organoids on day 7 of culture. **(D)** Percentage of CD40L^+^CD69^+^ cells of CD4^+^ T cells in ChAdOx1 nCoV-19-stimulated tonsil organoids with or without type 1 IFN or IL-6R blockade. **(E)** Percentage of CD4^+^ T cells co-expressing CD40L and cytokines (IFN-γ, TNF, and IL-2) in ChAdOx1 nCoV-19-stimulated tonsil organoids with or without type 1 IFN or IL-6R blockade. Data in A-E are combined from 2 experiments with a total of 8 donors. Each symbol represents an individual donor. Values in B and C were compared using Wilcoxon matched pairs signed rank test, values in D and E using Friedman test with Dunn’s multiple comparisons test.

Given that type I IFN and IL-6 modulated humoral responses, we sought to determine if these pathways also modulated the CD4^+^ T cell response. Although blocking type 1 IFN or IL-6 signaling did not affect the frequency of CD69^+^CD40L^+^ CD4^+^ T cells (Figure 6D), both treatments decreased cytokine production by CD4^+^ T cells in response to ChAdOx1 nCoV-19 (Figure 6E). This suggests that type 1 IFN and IL-6 both play a role in augmenting the CD4^+^ T cell effector response stimulated by ChAdOx1 nCoV-19.

One particularly important CD4^+^ T cell subset in promoting B cell responses are follicular helper T (T_FH_) cells, which regulate germinal center reactions (Crotty, 2014). We identified these cells by gating for PD-1^+^CXCR5^+^CD4^+^ T cells (Figure 7A). IL-6R blockade reduced the percentage of T_FH_ cells both in unstimulated and stimulated wells (Figure 7A-B). This finding is likely due to the requirement of IL-6 to maintain the T_FH_ phenotype (Korn and Hiltensperger, 2021). However, with regards to function, blocking IL-6 signaling only resulted in a modest decrease in TNF-producing T_FH_ and even a modest increase of CD40L^+^CD69^+^ T_FH_ cells, suggesting less of an effect on T_FH_ activation (Figure 7C-D). Conversely, blocking type 1 IFN did not affect the number of T_FH_ cells, but had a strong inhibitory effect on both IFN-γ and TNF production by T_FH_ cells. When examining activated T_FH_ as a percentage of total CD4^+^ T cells, both treatments resulted in decreased frequencies of cytokine-producing T_FH_ cells in the culture and no change in the percentage of CD40L^+^CD69^+^ T_FH_ cells (Figure 7E-F). Overall, these data show that IL-6 and type 1 IFN both modulate T_FH_ activation through different mechanisms: through maintaining T_FH_ phenotype and augmenting cytokine production, respectively.

**Figure 7:**
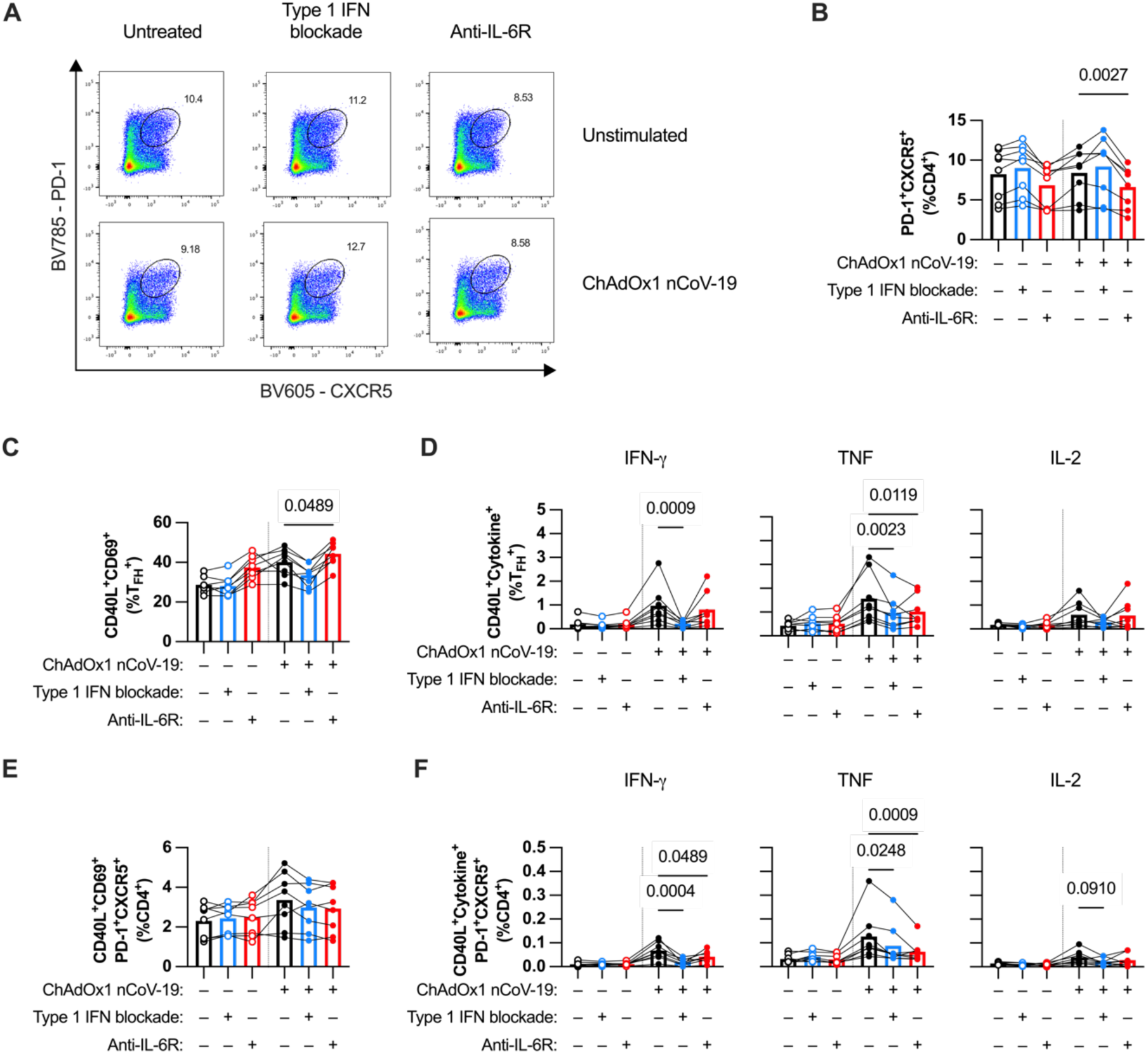
Type 1 IFN and IL-6 augment T follicular helper (T_FH_) cell responses to ChAdOx1 nCoV-19 through different mechanisms. **(A)** Representative FACS plots of T_FH_ (PD-1^+^CXCR5^+^) gating on CD4^+^ T cells in ChAdOx1 nCoV-19-stimulated tonsil organoids with or without type 1 IFN or IL-6R blockade. **(B)** Percentage of T_FH_ cells (of total CD4^+^ T cells) with or without type 1 IFN or IL-6R blockade. **(C)** Percentage of CD40L^+^CD69^+^ cells as a fraction of CD4^+^ T_FH_ cells in ChAdOx1 nCoV-19-stimulated tonsil organoids with or without type 1 IFN or IL-6R blockade. **(D)** Percentage of cells expressing both CD40L and cytokines (IFN-γ, TNF, and IL-2) as a fraction of CD4^+^ T_FH_ cells in ChAdOx1 nCoV-19-stimulated tonsil organoids with or without type 1 IFN or IL-6R blockade. **(E)** Percentage of CD40L^+^CD69^+^ CD4^+^ T_FH_ cells as a fraction of total CD4^+^ T cells in ChAdOx1 nCoV-19-stimulated tonsil organoids with or without type 1 IFN or IL-6R blockade. **(F)** Percentage of CD4^+^ T_FH_ cells expressing both CD40L and cytokines (IFN-γ, TNF, and IL-2) as a fraction of total CD4^+^ T cells in ChAdOx1 nCoV-19-stimulated tonsil organoids with or without type 1 IFN or IL-6R blockade. Data in B-F are combined from 2 experiments with a total of 8 donors. Each symbol represents an individual donor. Values in B-F were compared using Friedman test with Dunn’s multiple comparisons test.

## DISCUSSION

In this study we have validated the use of an in vitro human tonsil organoid system for studying the humoral responses to adenovirus vector vaccines. We could study vaccine antigen-specific antibody production and B and T cell activation, as well as bulk B cell activation, which allows the investigation of the mechanisms underlying these responses. Our findings demonstrate the central role of plasmacytoid dendritic cells (pDCs) in modulating humoral responses, primarily through type 1 IFN and indirectly through driving IL-6 signaling. Type 1 IFN and IL-6 also modulate CD4^+^ T cell responses, including T_FH_ cell responses, providing other potential mechanisms that regulate humoral responses to adenoviral vaccines. We summarize these findings in our model (Figure 8).

**Figure 8:**
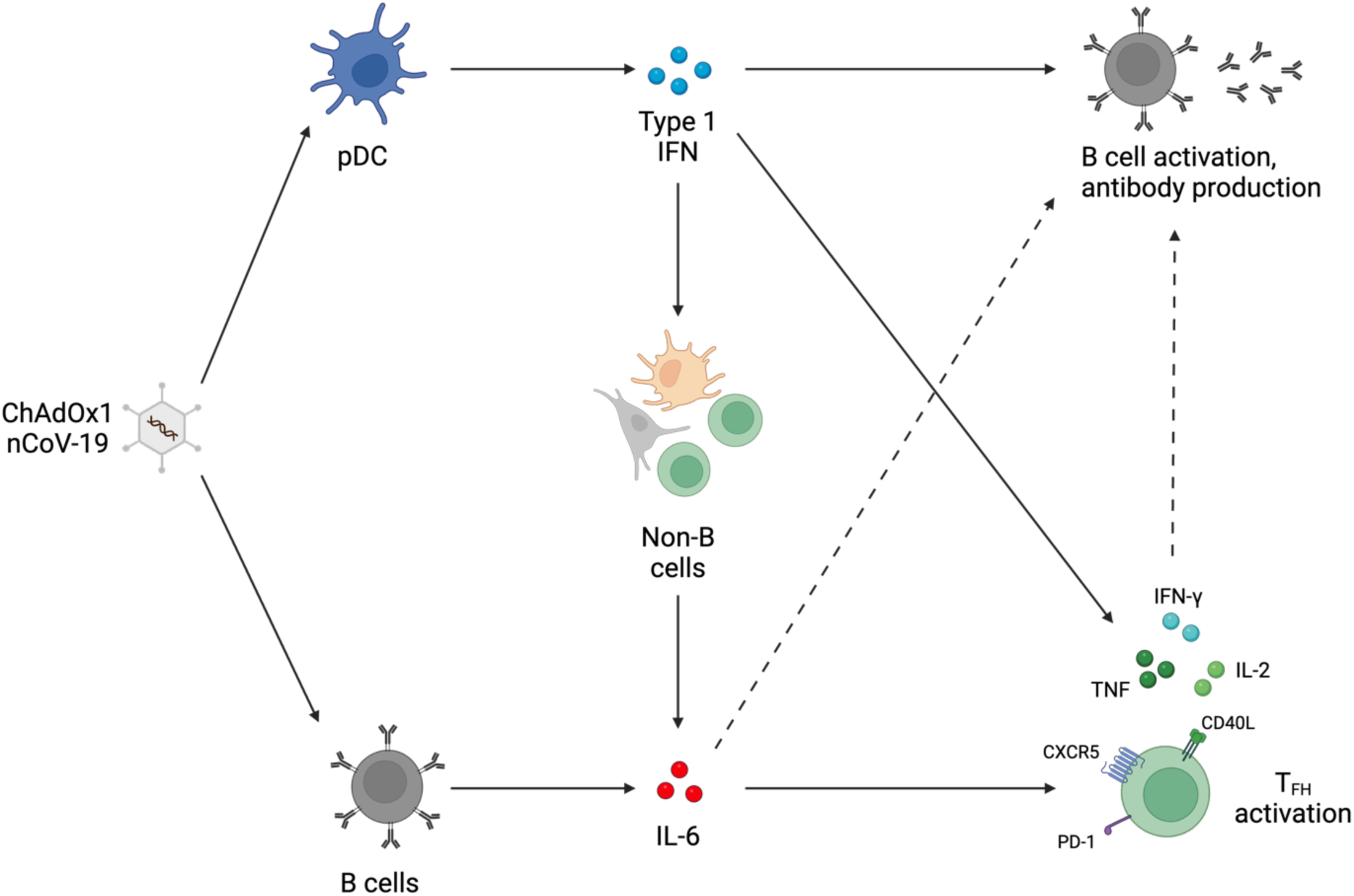

The in vitro tonsil organoid model has mainly been previously utilized for investigating responses to influenza vaccines (Kastenschmidt *et al*., 2023; Yin *et al*., 2023; Mitul *et al*., 2024). Although previous studies focused on influenza vaccines, it has been shown that adenoviral vectors can induce antibody responses in the model (Wagar *et al*., 2021). We investigated further the cell-cell interactions and signaling modulating this humoral response and extended these investigations into the regulation of CD4^+^ T cell, and specifically T_FH_ cell, responses. Through using the system, we could model adaptive immune responses to ChAdOx1 nCoV-19 within lymphoid tissue and probe into mechanisms regulating these responses. Through combining cell depletions and cytokine blockade/supplementation, we could investigate intricate signaling interplay between different cells and cytokines, which revealed a critical pDC–IFN-α–IL-6 signaling axis. We believe this robustly validates the use of this model for studying entirely new classes of vaccines.

An important consideration with experimental models is identifying the aspects of the actual process that it captures. Our culture system appears to efficiently model a recall/boost response to adenoviral vectors rather than a “prime” response given that reliable detection of specific IgG antibody production requires pre-existing S1-specific B cells. However, IgM production could be measured but was stochastic likely due to limited frequency of naive B cells in the culture well. As such, our system appears to more robustly model a booster vaccination response in a previously exposed individual rather than a priming vaccination. The initial paper published on the model demonstrated priming of naive B cell responses either by using larger culture wells or specifically enriching for naive B cells, increasing the overall number of naïve B cells in the system as well as the chances of having antigen-specific naïve B cells in the culture (Wagar *et al*., 2021). These parameters should be considered when designing and implementing this organoid model.

With our system, we discovered that pDCs are central modulators of humoral responses to ChAdOx1 nCoV-19. The role of pDCs has rarely been discussed in the context of humoral responses to adenovirus vector responses. A clinical study reported that the number of pDCs in blood are negatively correlated with antibody responses to ChAdOx1 nCoV-19 (Ryan *et al*., 2023). Elevated pDC frequency in the blood might indicate lower recruitment into lymph nodes after vaccination, supporting the role of pDCs in adenovirus vector humoral responses generated in SLOs. A previous study using the tonsil organoid model also reported a sharp decrease in humoral responses towards LAIV with pDC-depletion (Wagar *et al*., 2021), suggesting that pDCs may also play a role in responses to multiple vaccine platforms.

Our findings show type 1 IFN as the main mediator of pDCs’ effect on humoral responses. A study in NHPs previously showed a correlation between type 1 IFN and spike-specific humoral responses induced by Ad26.COV2.S (He *et al*., 2022), and we have now directly demonstrated a mechanistic link with ChAdOx1 nCoV-19. Previous studies testing an adenovirus vector vaccine against foot and mouth disease virus (FMDV) in mice reported higher humoral and cellular responses when including porcine IFN-α in the construct (Su *et al*., 2013; Duan *et al*., 2020). Type 1 IFNs are known to support B cell activation and differentiation into plasma cells (Bekeredjian-Ding *et al*., 2005; Liu *et al*., 2019). Previous studies also report the role of type 1 IFN in augmenting humoral responses to other vaccine modalities including LAIV and MVA (Jego *et al*., 2003; Aljurayyan *et al*., 2016; Zhong *et al*., 2021), suggesting that this effect applies for humoral responses in general.

In addition to directly activating B cells, type 1 IFN also augmented humoral responses through promoting IL-6 secretion. In vivo studies in mice and hepatitis C patients receiving recombinant pegylated IFN-α injections show that type 1 IFN enhances production of IL-6 (Ito *et al*., 1996; Murray *et al*., 2015). Historically named B-cell stimulatory factor-2, IL-6 is known to promote survival, expansion, and maturation of B cells to become plasmablasts (Rose-John, Winthrop and Calabrese, 2017). This is supported by several clinical studies reporting decreased humoral responses to mRNA-based SARS-CoV-2 vaccines with anti-IL-6R treatment (Tobudic *et al*., 2023; Schiff *et al*., 2024).

Type 1 IFN and IL-6 also modulate bulk CD4^+^ T cell and T_FH_ responses in our experimental model. Type 1 IFN promotes Bcl6 expression in CD4^+^ T cells (Nakayamada *et al*., 2014), but is inadequate to induce complete T_FH_ differentiation. Full T_FH_ commitment requires other cytokines, including IL-6 (Harker *et al*., 2011). As type 1 IFN itself augments IL-6 secretion, it appears that type 1 IFN and IL-6 synergize to induce T_FH_ differentiation and activation in response to adenovirus vaccination, in turn promoting T-dependent B cell responses. Other than direct effect on T cells, type 1 IFN may also modulate cDCs to promote T_FH_ differentiation (Dahlgren *et al*., 2022). Studies of adenoviral FMDV vaccines in mice reported that including type 1 IFN in the construct enhanced both induction and activation of T_FH_ cells as well as the antibody production against the transgene (Su *et al*., 2013; Duan *et al*., 2020).

Besides their cytokine production, pDCs may also contribute to humoral responses through producing target antigens. Given their efficient transduction by ChAdOx1, it is possible that pDCs can present antigens to T cells or provide antigens to B cells for the GC reaction. Several studies have reported that pDC subsets can contribute to antigen presentation to T cells, either by directly presenting (Zhang *et al*., 2017; Alculumbre *et al*., 2018) or by transferring antigens to cDCs (Rogers *et al*., 2017; Fu *et al*., 2020). Our data suggests that their production of type 1 IFN is their major mode of action in this model, given the near complete restoration of antibody production in pDC-depleted cultures by recombinant IFN-α supplementation. Nonetheless, the decrease in antibody production with type 1 IFN blockade was milder than pDC-depletion, which suggests that antigen production may still form a minor part of pDCs’ role in the humoral response to adenovirus vectors.

The tonsil organoid system provides an in vitro model for studying vaccine responses with cells derived from human SLO tissue that resembles lymph nodes closer than peripheral blood culture models. However, as specialized SLOs that are constantly exposed to pathogens in the oral tract, tonsils would still have differences with lymph nodes, and this unique biology may be particularly relevant when considering mucosal vaccination. The pathology prompting the removal of the tonsils may also affect the responses in the system, as evidenced by several donors that produced S1-specific antibody responses at baseline (and were thus excluded from analysis).

Using the in vitro tonsil organoid model, we were able to demonstrate humoral and cellular responses to ChAdOx1 nCoV-19. The system appears to model recall responses, which would be applicable to booster vaccinations. We discovered that pDCs modulate humoral responses to adenoviral vector vaccines, primarily through type 1 IFN. Type 1 IFN augmented IL-6 production, which also modulated humoral responses. Both type 1 IFN and IL-6 promoted CD4^+^ T and T_FH_ responses to the transgene, possibly partially linking cellular and humoral immunity. The system provides great potential for mechanistic studies on vaccine responses.

## METHODS

### Tonsil tissue collection & processing

Whole human tonsils were collected from January 2022 to October 2023 from adult or pediatric patients undergoing tonsillectomy for sleep apnea or any other indication for elective tonsillectomy. Patients receiving immunosuppressive treatment, with severe inflammatory or exudative lesions on the tonsils, or with underlying immunocompromising disease were excluded. Ethics approval was obtained from the Oxford Radcliffe Biobank research tissue bank ethics (reference 19/SC/0173).

Tonsil tissue was processed as previously described (Wagar *et al*., 2021). Whole tonsils were collected in phosphate buffer saline (PBS) after surgery, then decontaminated by immersion in antimicrobial media (PBS, 1% penicillin, 1% streptomycin, 0.01% Normocin (Invivogen)) for ≥1 hour at 4°C. Tonsils were dissected and ground through a 70-micron strainer to obtain single-cell suspensions. To decrease cell debris, cells were isolated by Ficoll density gradient separation and washed with complete medium (RPMI-1640 with glutamine, 10% fetal bovine serum (FBS), 1% nonessential amino acids, 0.01% Normocin, 1% insulin/transferrin/selenium). Cells were counted, frozen in FBS with 10% dimethyl sulfoxide (DMSO), and stored at −150°C.

Pre-pandemic tonsil samples were collected prior to 2019 under the GI Biobank ethics (reference: 16/YH/0247) and processed and stored as previously described (Hagel *et al*., 2021). Written informed consent was received from all tissue donors, or their respective legal guardians, as appropriate. All work was performed in compliance with the principles of the Declaration of Helsinki (2008) and in accordance with relevant ethical regulation.

### Tonsil organoid culture

Tonsil organoid cultures were performed as previously described (Wagar *et al*., 2021). Cell aliquots were thawed and resuspended in complete media at 4×10^7^ cells/mL. Cells were plated in 96-well permeable (0.4-μm pore size) transwell polycarbonate membrane plates (Corning), with the upper chamber containing 1×10^6^ or2×10^6^ cells, depending on the experiment, and the lower chamber containing media. Culture media was supplemented with 1 μg ml^−1^ recombinant human B cell-activating factor (BAFF; BioLegend). Cultures were stimulated using ChAdOx1 nCoV-19 (multiplicity of infection (MOI): 1000 viral particles/cell). ChAdOx1 nCoV-19 was used either as discarded clinical product (OUH Pharmacy) or produced by the Pandemic Sciences Institute (University of Oxford) Viral Vector Core Facility. Cultures were incubated at 37°C and 5% CO_2_. Culture media in the bottom chamber was replaced with fresh media every three days.

pDC-depletion was done using magnetic separation using the CD303 (BDCA-2) MicroBead Kit, per manufacturer’s instructions (Miltenyi Biotec). Tonsil cells were incubated with biotinylated anti-BDCA-2 antibodies and subsequently with anti-biotin magnetic beads. Cells were then passed through LS separation columns (Miltenyi Biotec) placed on magnetic stands. pDC-depleted cells were collected from the flowthrough and cultured as described above. IFN-α and IL-6R blockade treatment used B18R (eBioscience, final concentration: 10 μg/mL) and anti-human IL-6R (InVivoSim, Bio X Cell, final concentration: 10 μg/mL). IFN-α supplementation used IFN-α (Sigma-Aldrich) with a final concentration of 50 ng/ml.

B cell separation was done using magnetic separation using the CD19 Microbead Kit, per manufacturer’s instructions (Miltenyi Biotec). Tonsil cells were incubated with magnetic beads conjugated to anti-CD19 antibodies. Cells were then passed through LD separation columns (Miltenyi Biotec) placed on magnetic stands. B cell depleted cells were collected from the flowthrough, while selected B cells were collected by flushing the columns after removing LD columns from the magnetic stands. B cell depleted cells and selected B cells were then cultured as described above.

### Specific antibody measurement

Culture supernatants were harvested from the lower chamber of the transwell plates, supernatants were collected every three days starting from day 6 of culture alongside the media change. Specific IgG and IgM antibody levels were measured in culture supernatants by ELISA using LEGEND MAX^TM^ SARS-CoV-2 Spike S1 Human IgG ELISA kit (BioLegend) and LEGEND MAX^TM^ SARS-CoV-2 Spike S1 Human IgM ELISA kit (BioLegend), respectively, per manufacturer’s instructions. Samples with calculated antibody titers below the concentration of the lowest standard were converted to half of the lowest standard.

### Generation of B cell tetramers

Antigen-specific B cells were identified using binding to fluorescently labelled SARS-CoV-2 spike S1 region tetramers. S1 tetramers were generated as previously described (Oberhardt *et al*., 2021). Tetramerization was done through addition of PE- or BV421-conjugated streptavidin (Biolegend) to biotinylated S1 proteins (Biolegend). Streptavidin was added incrementally at one-fifth of the amount of S1 protein, which was repeated five times with 20 minutes incubation at 4°C in between.

### Flow cytometry

Tonsil organoid cells were harvested from the upper chamber by washing with PBS. Cells were then washed twice in FACS buffer (PBS + 0.05% BSA + 2mM EDTA) before being incubated with the surface staining cocktail for 30 minutes light-protected at 4°C. Cells were washed twice in FACS buffer, fixed using BD Cytofix (BD Biosciences) or BD Cytofix/Cytoperm (BD Biosciences) for 20 minutes at 4°C, and washed twice with FACS buffer. Samples were stored at 4°C before running on the flow cytometer. Flow cytometry was done using BD LSRFortessa Cell Analyzer or BD LSR II Flow Cytometer (BD Biosciences).

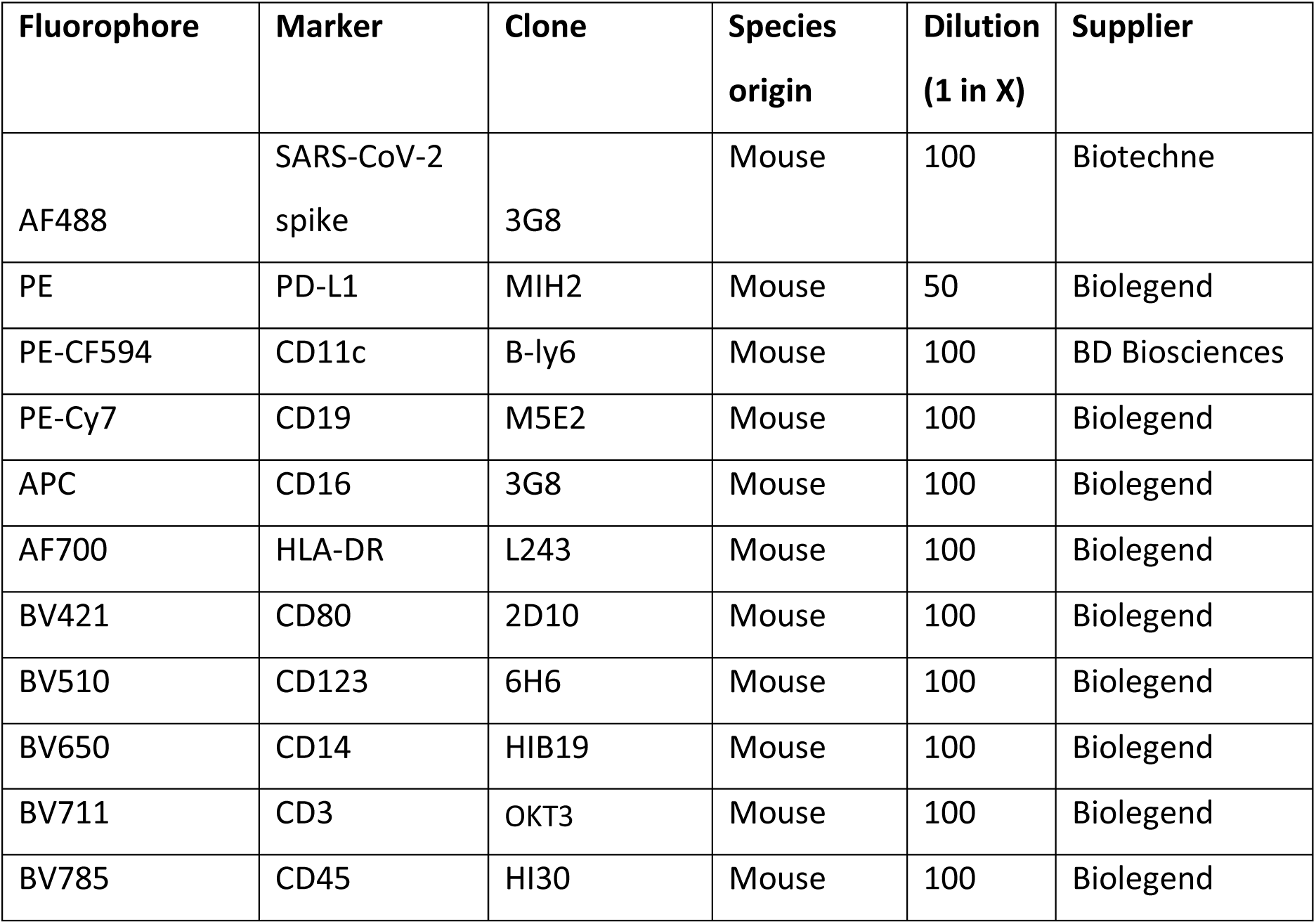
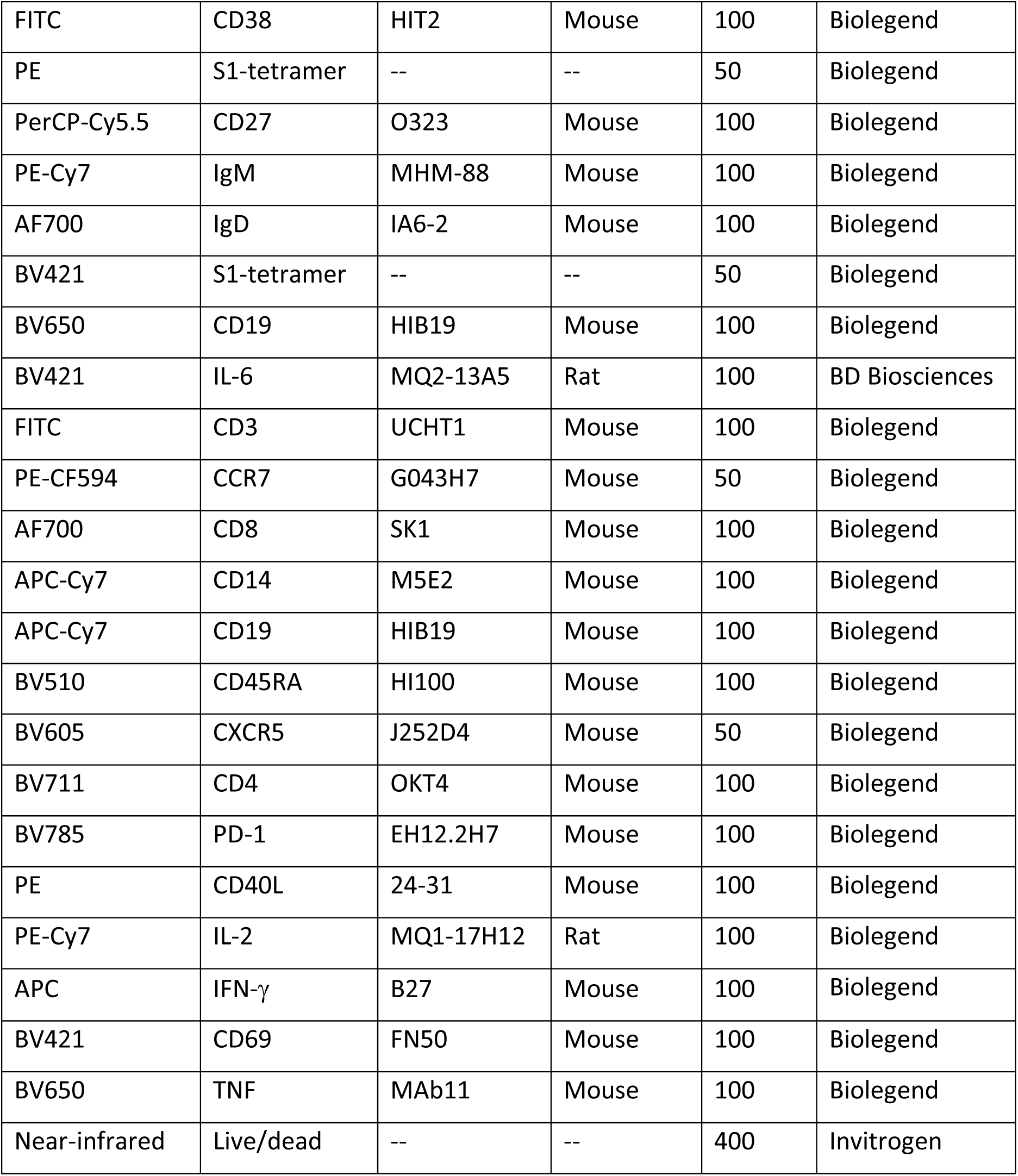

### T cell assays

For T cell assays, tonsil organoids were plated as described above and cultured up to 7 days. Brefeldin A was added to cultures at 16-20 hours before harvest to enhance intracellular cytokine detection. Cells were harvested, washed, and incubated in the surface staining cocktail as described above. Cells were then fixed and permeabilized by being incubated in BD Cytofix/Cytoperm (BD Biosciences) for 20 minutes at 4°C. Fixed cells were incubated with the intracellular staining cocktail for 30 minutes at 4°C and washed twice before running the samples in the flow cytometer. Flow cytometry was done using BD LSRFortessa Cell Analyzer or BD LSR II Flow Cytometer (BD Biosciences).

### Cytokine measurements

Cytokine levels were measured from culture media supernatants harvested 24 hours after plating using LegendPlex^TM^ Human Inflammation Panel 1 (Biolegend), per manufacturer’s instructions. Briefly, supernatants were incubated with detection beads coated with antibodies against the panel cytokines for two hours. Beads were then incubated with biotinylated detection antibodies and subsequently with PE-conjugated streptavidin. Beads are then detected through flow cytometry (BD LSRFortessa Cell Analyzer), then converted back to cytokine concentrations using Qognit software (Biolegend).

### Data analysis

Analysis of flow cytometry data was performed using FlowJo version 10.8.1. Statistical analyses were performed in GraphPad Prism version 9. Comparison of different donor groups (Figure 1G) were done using a nonparametric Mann-Whitney U test. Paired samples (comparison of different treatments for matched/pairs donors) were done using Wilcoxon matched pairs signed rank test (for two groups) or using Friedman test with Dunn’s multiple comparisons test (for three groups). For figures 5F-G, comparisons were only made between unstimulated and stimulated groups separately for each treatment. For figures 6D-E and 7B-F, comparisons were only made between stimulated groups with different treatments. Correlation tests were done using Spearman test.

## ACKNOWLEDGEMENTS

We thank the patients who consented to donate their tissue for the research, as well as the Oxford Radcliffe Biobank for patient consenting and sample collection. We thank Helen Ferry for technical assistance in flow cytometry. We thank Marta Rizzi (University of Freiburg) for assistance in generating the SARS-CoV-2 spike peptide tetramers.

## AUTHOR CONTRIBUTIONS

Conceptualization: N.M.P.

Methodology: M.F.P. and N.M.P.

Formal Analysis: M.F.P.

Investigation: M.F.P.

Sample provision: M.F.P. and K.P.

Reagent provision: E.B.

Data Curation: M.F.P.

Writing – Original Draft: M.F.P.

Writing – Review & Editing: M.F.P., K.P., E.B., P.K., and N.M.P. Visualization: M.F.P.

Supervision: P.K. and N.M.P.

Funding Acquisition: P.K. and N.M.P.

---

**Figure S1:**
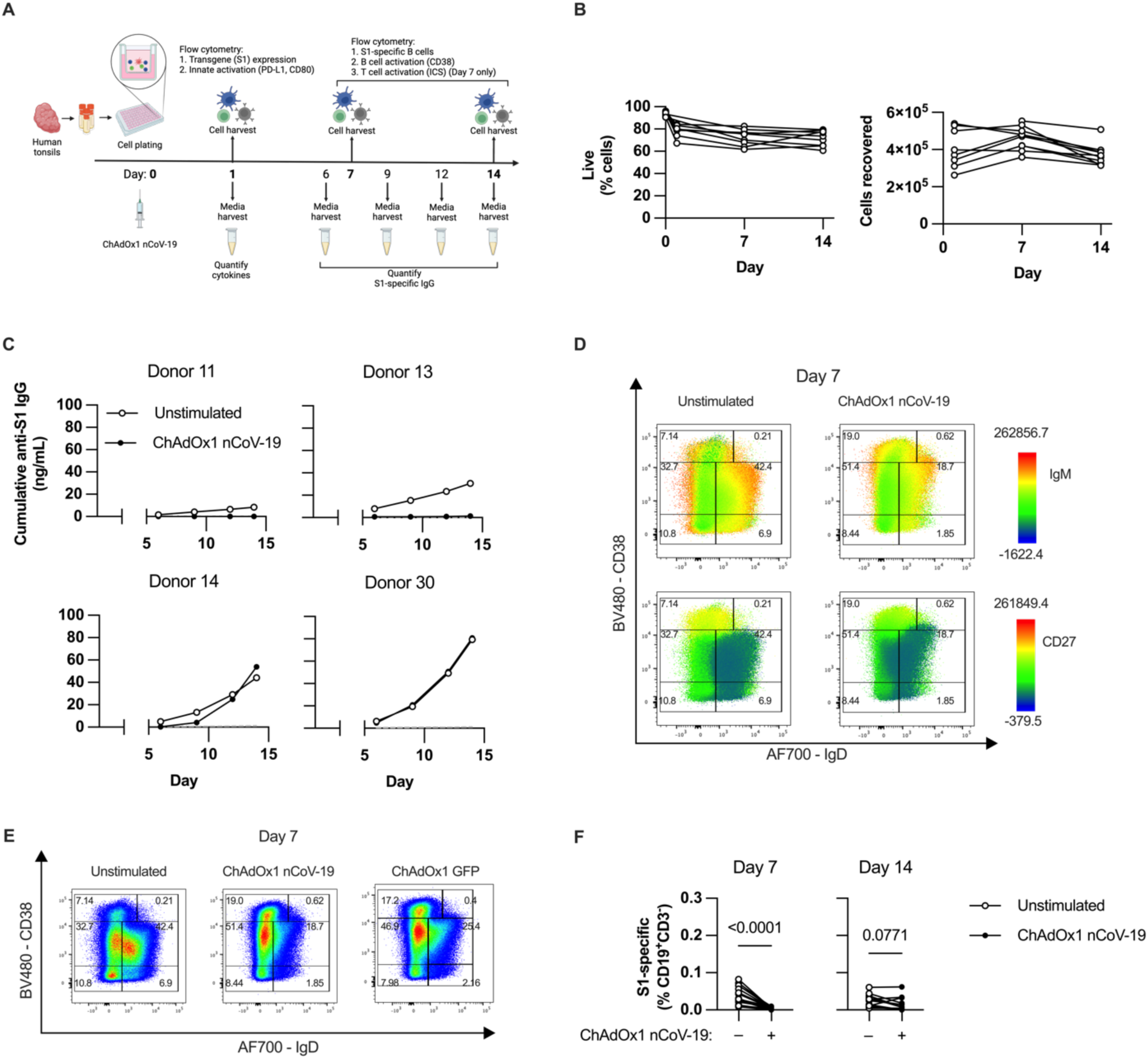
**(A)** Experimental schematic. **(B)** Viability and cell recovery counts across donors in 14-day organoid cultures with the initial plating of 2E6 cells/well. **(C)** Several post-pandemic tonsil donors produce anti-S1 IgG at baseline, which is initially suppressed with ChAdOx1 nCoV-19 stimulation. These donors were removed from subsequent analyses. **(D)** Representative FACS plot of IgM and CD27 expression (overlaid on IgD and CD38) on B cells (singlet, live, CD45^+^CD19^+^CD3^-^) at day 7 of organoid culture. **(E)** Representative flow plot of B cells on day 7 of organoid culture with ChAdOx1 nCoV-19 and ChAdOx1 GFP stimulation. **(F)** Percentage of S1-specific B cells in post-pandemic donors with/without ChAdOx1 nCoV-19 stimulation at day 7 and 14 of culture. Data in F is combined from 4 experiments with a total of 16 donors. Each symbol represents an individual donor. Values in F were compared using Wilcoxon matched pairs signed rank test.

**Figure S2:**
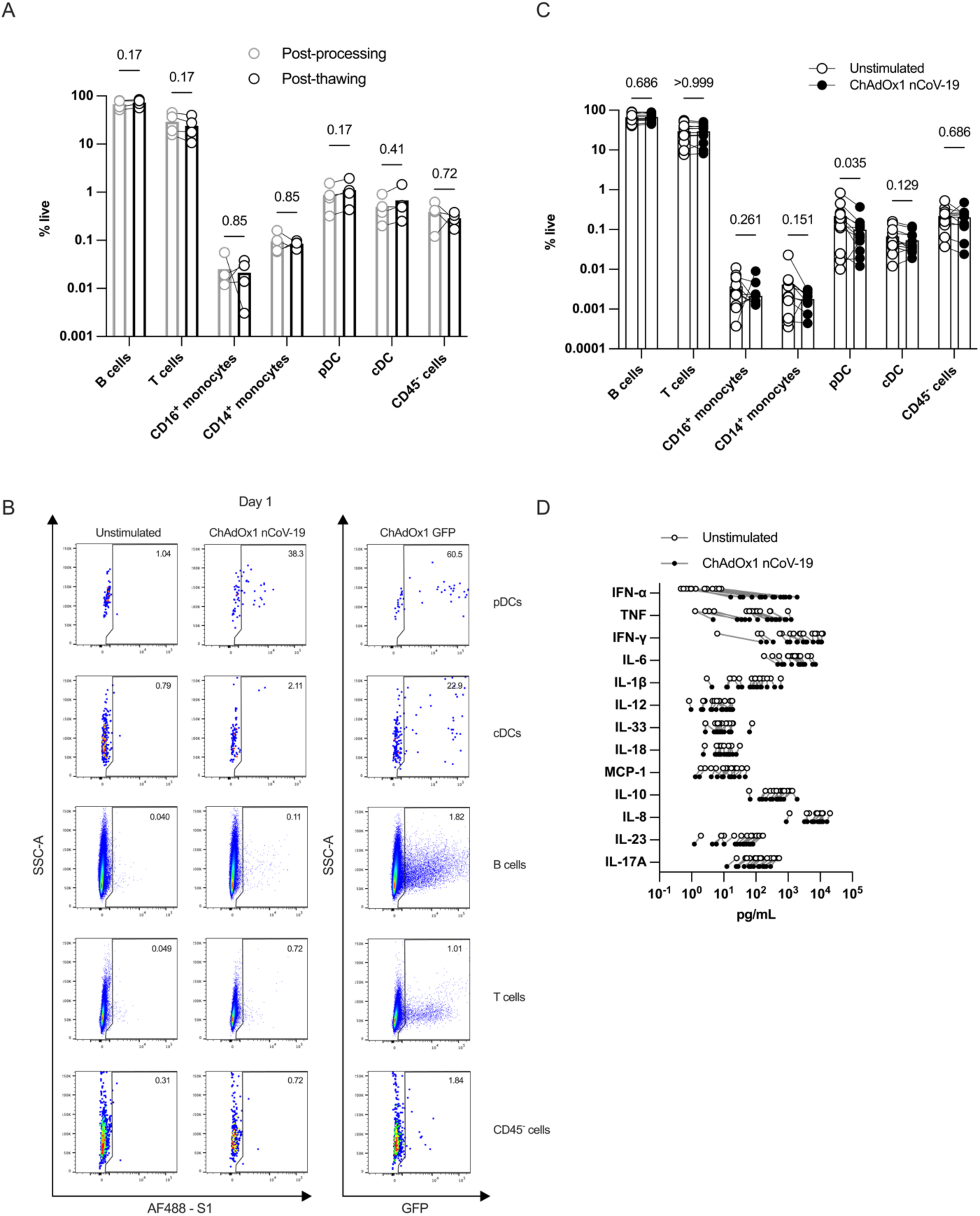
**(A)** Cell composition of tonsil cells after processing fresh tissue and directly after thawing cryopreserved cells (from 4 donors). **(B)** Representative FACS plots of transduction of different cell types based on detection of SARS-CoV-2 S1 or GFP. **(C)** Cell composition of tonsil cells after 24-hour cultures with or without ChAdOx1 nCoV-19 stimulation. **(D)** Individual cytokine levels for tonsil organoid supernatants after 24-hour cultures with or without ChAdOx1 nCoV-19 stimulation. Data in A are from 4 donors, data in C are combined from 4 experiments with a total of 15 donors, data in D are combined from 4 experiments with a total of 16 donors. Each symbol represents an individual donor. Values in A were compared using multiple paired t-tests.

**Figure S3:**
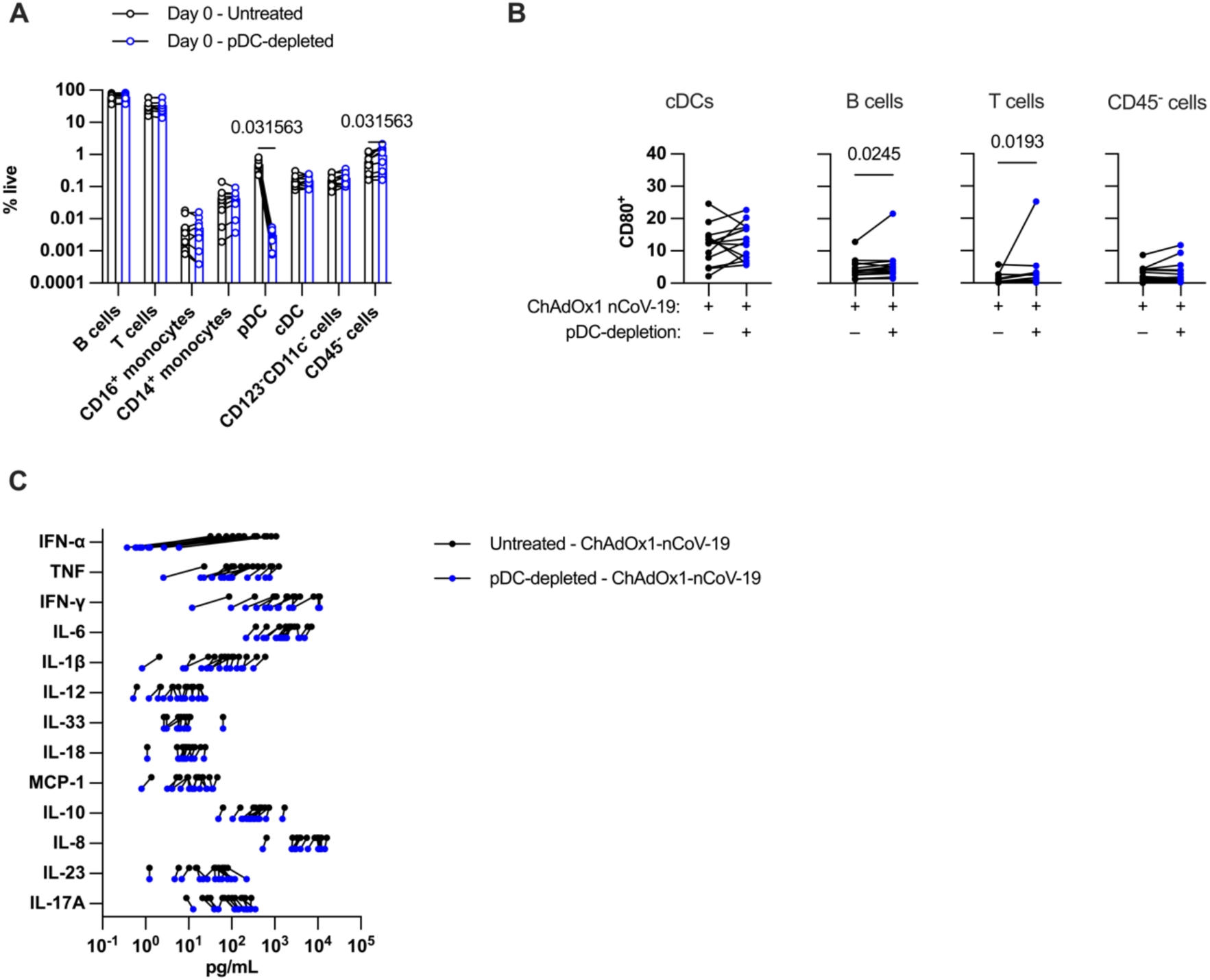
**(A)** Composition of tonsil cell frequency before and after pDC-depletion. pDC-depletion was carried out using magnetic bead separation against BDCA2. **(B)** Surface expression of CD80 for different cell types in ChAdOx1 nCoV-19-stimulated organoids with or without pDC-depletion. **(C)** Individual cytokine levels for tonsil organoid supernatants after 24-hour ChAdOx1 nCoV-19 stimulation in cultures with or without pDC-depletion. Data in A-C are combined from 4 experiments with a total of 14 donors. Each symbol represents an individual donor. Values in A and B were compared using multiple Wilcoxon matched pairs signed rank test.

**Figure S4:**
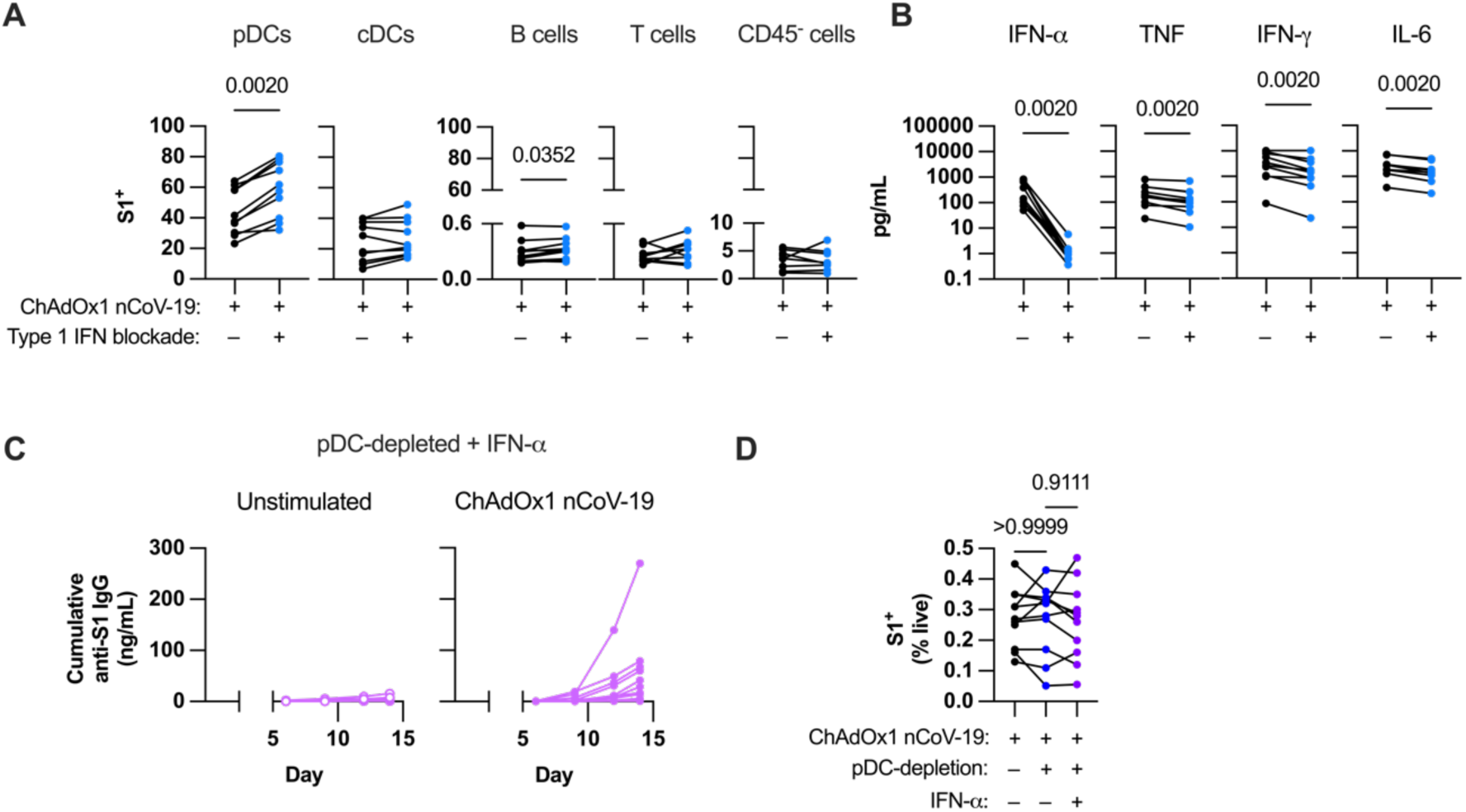
**(A)** Cytokine levels in ChAdOx1 nCoV-19-stimulated tonsil organoid supernatants with or without type 1 IFN blockade. **(B)** Transduction rates of different cell types in ChAdOx1 nCoV-19-stimulated organoids with or without type 1 IFN blockade. **(C)** Cumulative anti-S1 IgG production over time in pDC-depleted ChAdOx1 nCoV-19-stimulated tonsil organoids with IFN-α supplementation. **(D)** Percentage of total transduced cells after 24-hour culture in pDC-depleted ChAdOx1 nCoV-19-stimulated organoids and with or without IFN-α supplementation. Data in A-B are combined from 3 experiments with a total of 10 donors, C-D from 3 experiments with 11 donors. Each symbol represents an individual donor. Values in A-B were compared using Wilcoxon matched pairs signed rank test, values in D using Friedman test with Dunn’s multiple comparisons test.

**Figure S5:**
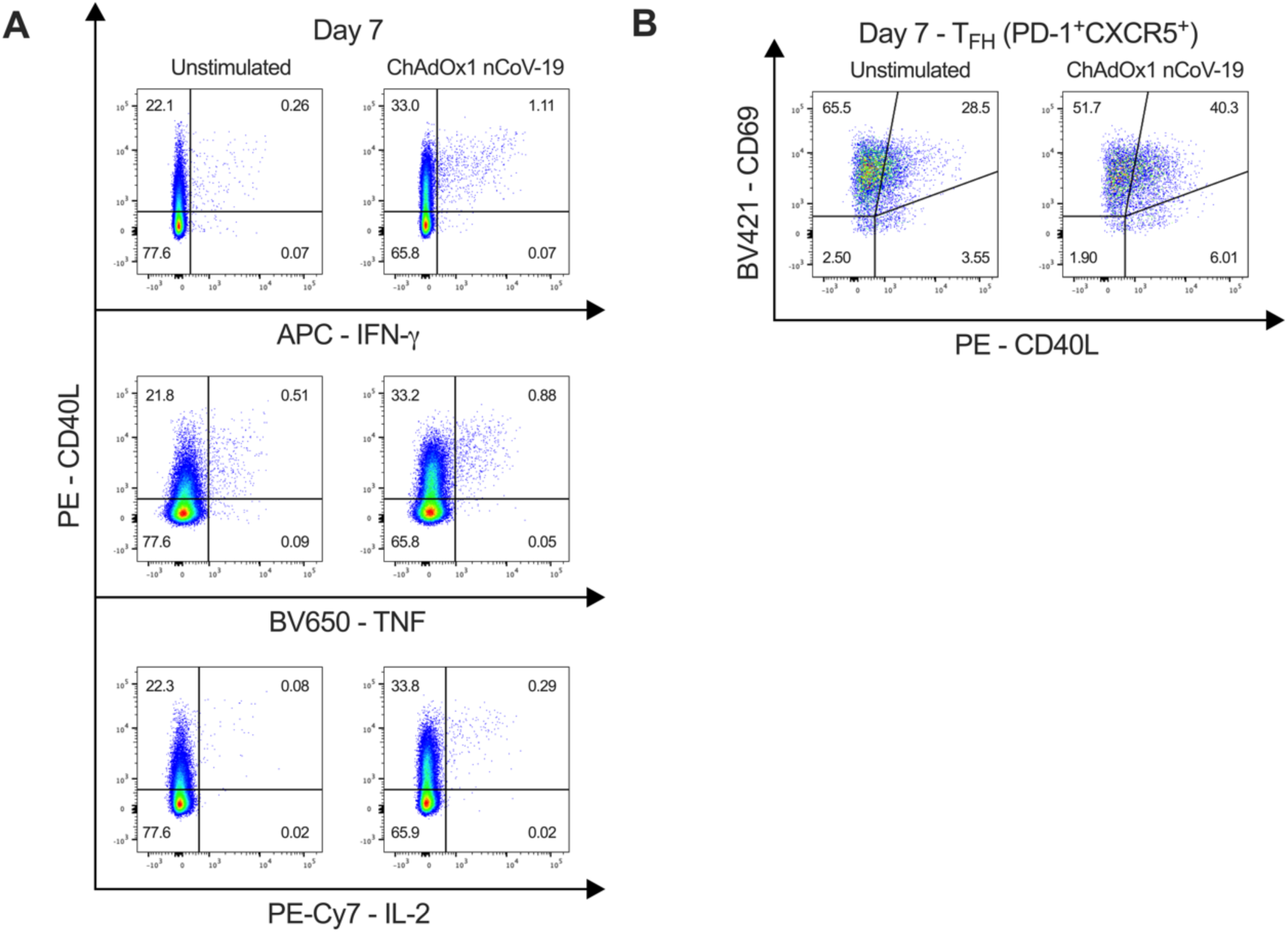
**(A)** Representative FACS plots of CD40L against cytokines (IFN-γ, TNF, and IL-2) for CD4 T cells in tonsil organoids. (B) Representative FACS plots of CD40L and CD69 expression on PD-1^+^CXCR5^+^ (T_FH_) cells in tonsil organoids.

## REFERENCES

Aarden, L.A. et al. (1987) ‘Production of hybridoma growth factor by human monocytes’, European Journal of Immunology, 17(10), pp. 1411–1416. 10.1002/eji.1830171004.

Alculumbre, S.G. et al. (2018) ‘Diversification of human plasmacytoid predendritic cells in response to a single stimulus’, Nature Immunology, 19(1), pp. 63–75. 10.1038/s41590-017-0012-z.

Aljurayyan, A.N. et al. (2016) ‘A critical role of T follicular helper cells in human mucosal anti-influenza response that can be enhanced by immunological adjuvant CpG-DNA’, Antiviral Research, 132, pp. 122–130. 10.1016/j.antiviral.2016.05.021.

Bekeredjian-Ding, I.B. et al. (2005) ‘Plasmacytoid dendritic cells control TLR7 sensitivity of naive B cells via type I IFN’, The Journal of Immunology, 174(7), pp. 4043–4050. 10.4049/jimmunol.174.7.4043.

Crotty, S. (2014) ‘T follicular helper cell differentiation, function, and roles in disease’, Immunity, 41(4), pp. 529–542. 10.1016/j.immuni.2014.10.004.

Dahlgren, M.W. et al. (2022) ‘Type I interferons promote germinal centers through B cell intrinsic signaling and dendritic cell dependent Th1 and Tfh cell lineages’, Frontiers in Immunology, 13, p. 932388. 10.3389/fimmu.2022.932388.

Duan, X. et al. (2020) ‘1IFN-α modulates memory Tfh cells and memory B cells in mice, following recombinant FMDV adenoviral challenge’, Frontiers in Immunology, 11, p. 701. 10.3389/fimmu.2020.00701.

Feldman, J. et al. (2021) ‘Naive human B cells engage the receptor binding domain of SARS-CoV-2, variants of concern, and related sarbecoviruses’, Science Immunology, 6(66), p. eabl5842. 10.1126/sciimmunol.abl5842.

Foster, W.S. et al. (2022) ‘Tfh cells and the germinal center are required for memory B cell formation & humoral immunity after ChAdOx1 nCoV-19 vaccination’, Cell Reports Medicine, 3(12), p. 100845. 10.1016/j.xcrm.2022.100845.

Fourati, S. et al. (2022) ‘Pan-vaccine analysis reveals innate immune endotypes predictive of antibody responses to vaccination’, Nature Immunology. 10.1038/s41590-022-01329-5.

Fu, C. et al. (2020) ‘Plasmacytoid dendritic cells cross-prime naive CD8 T cells by transferring antigen to conventional dendritic cells through exosomes’, Proceedings of the National Academy of Sciences, 117(38), pp. 23730–23741. 10.1073/pnas.2002345117.

Hagan, T. et al. (2022) ‘Transcriptional atlas of the human immune response to 13 vaccines reveals a common predictor of vaccine-induced antibody responses’, Nature Immunology. 10.1038/s41590-022-01328-6.

Hagel, J.P. et al. (2021) ‘Defining T cell subsets in human tonsils using chipcytometry’, The Journal of Immunology, 206(12), pp. 3073–3082. 10.4049/jimmunol.2100063.

Halperin, S.A. et al. (2022) ‘Final efficacy analysis, interim safety analysis, and immunogenicity of a single dose of recombinant novel coronavirus vaccine (adenovirus type 5 vector) in adults 18 years and older: an international, multicentre, randomised, double-blinded, placebo-controlled phase 3 trial’, The Lancet, 399(10321), pp. 237–248. 10.1016/S0140-6736(21)02753-7.

Harker, J.A. et al. (2011) ‘Late interleukin-6 escalates T follicular helper cell responses and controls a chronic viral infection’, Science, 334(6057), pp. 825–829. 10.1126/science.1208421.

He, X. et al. (2022) ‘A homologous or variant booster vaccine after Ad26.COV2.S immunization enhances SARS-CoV-2–specific immune responses in rhesus macaques’, Science Translational Medicine, 14(638), p. eabm4996. 10.1126/scitranslmed.abm4996.

Ito, N. et al. (1996) ‘Induction of interleukin-6 by interferon alfa and its abrogation by a serine protease inhibitor in patients with chronic hepatitis C’, Hepatology, 23(4), pp. 669–675. 10.1002/hep.510230403.

Jego, G. et al. (2003) ‘Plasmacytoid dendritic cells induce plasma cell differentiation through type I interferon and interleukin 6’, Immunity, 19(2), pp. 225–234. 10.1016/S1074-7613(03)00208-5.

Jiang, M. et al. (2023) ‘COVID-19 adenovirus vector vaccine induces higher interferon and pro-inflammatory responses than mRNA vaccines in human PBMCs, macrophages and moDCs’, Vaccine, 41(26), pp. 3813–3823. 10.1016/j.vaccine.2023.04.049.

Kastenschmidt, J.M. et al. (2023) ‘Influenza vaccine format mediates distinct cellular and antibody responses in human immune organoids’, Immunity, 56, pp. 1–17. 10.1016/j.immuni.2023.06.019.

Korn, T. and Hiltensperger, M. (2021) ‘Role of IL-6 in the commitment of T cell subsets’, Cytokine, 146, p. 155654. 10.1016/j.cyto.2021.155654.

Li, R. et al. (2018) ‘Toll-like receptor 4 signalling regulates antibody response to adenoviral vector-based vaccines by imprinting germinal centre quality’, Immunology, 155(2), pp. 251–262. 10.1111/imm.12957.

Liu, M. et al. (2019) ‘Type I interferons promote the survival and proinflammatory properties of transitional B cells in systemic lupus erythematosus patients’, Cellular & Molecular Immunology, 16(4), pp. 367–379. 10.1038/s41423-018-0010-6.

Mitul, M.T. et al. (2024) ‘Tissue-specific sex differences in pediatric and adult immune cell composition and function’, Frontiers in Immunology, 15, p. 1373537. 10.3389/fimmu.2024.1373537.

Murray, C. et al. (2015) ‘Interdependent and independent roles of type I interferons and IL-6 in innate immune, neuroinflammatory and sickness behaviour responses to systemic poly I:C’, Brain, Behavior, and Immunity, 48, pp. 274–286. 10.1016/j.bbi.2015.04.009.

Nakayamada, S. et al. (2014) ‘Type I IFN induces binding of STAT1 to Bcl6: divergent roles of STAT family transcription factors in the T follicular helper cell genetic program’, The Journal of Immunology, 192(5), pp. 2156–2166. 10.4049/jimmunol.1300675.

Oberhardt, V. et al. (2021) ‘Rapid and stable mobilization of CD8+ T cells by SARS-CoV-2 mRNA vaccine’, Nature, 597(7875), pp. 268–273. 10.1038/s41586-021-03841-4.

Provine, N.M. et al. (2016) ‘Transient CD4^+^ T cell depletion results in delayed development of functional vaccine-elicited antibody responses’, Journal of Virology. Edited by F. Kirchhoff, 90(9), pp. 4278–4288. 10.1128/JVI.00039-16.

Provine, N.M. et al. (2021) ‘MAIT cell activation augments adenovirus vector vaccine immunogenicity’, Science, 371(6528), pp. 521–526. 10.1126/science.aax8819.

Provine, N.M. and Klenerman, P. (2022) ‘Adenovirus vector and mRNA vaccines: Mechanisms regulating their immunogenicity’, European Journal of Immunology, p. eji.202250022. 10.1002/eji.202250022.

Rogers, G.L. et al. (2017) ‘Plasmacytoid and conventional dendritic cells cooperate in crosspriming AAV capsid-specific CD8+ T cells’, Blood, 129(24), pp. 3184–3195. 10.1182/blood-2016-11-751040.

Rose-John, S., Winthrop, K. and Calabrese, L. (2017) ‘The role of IL-6 in host defence against infections: immunobiology and clinical implications’, Nature Reviews Rheumatology, 13(7), pp. 399–409. 10.1038/nrrheum.2017.83.

Ryan, F.J. et al. (2023) ‘A systems immunology study comparing innate and adaptive immune responses in adults to COVID-19 mRNA and adenovirus vectored vaccines’, Cell Reports Medicine, 4(3), p. 100971. 10.1016/j.xcrm.2023.100971.

Sadoff, J. et al. (2022) ‘Final analysis of efficacy and safety of single-dose Ad26.COV2.S’, New England Journal of Medicine, 386(9), pp. 847–860. 10.1056/NEJMoa2117608.

Sanz, I. et al. (2019) ‘Challenges and opportunities for consistent classification of human B cell and plasma cell populations’, Frontiers in Immunology, 10, p. 2458. 10.3389/fimmu.2019.02458.

Schiff, A.E. et al. (2024) ‘Immunomodulators and risk for breakthrough COVID-19 after third SARS-CoV-2 mRNA vaccine among patients with rheumatoid arthritis: a cohort study’, Annals of the Rheumatic Diseases, 83(5), pp. 680–682. 10.1136/ard-2023-225162.

Su, C., et al. (2013) ‘IFN-α as an adjuvant for adenovirus-vectored FMDV subunit vaccine through improving the generation of T follicular helper cells’, PLoS ONE. Edited by M.M. Rodrigues, 8(6), p. e66134. 10.1371/journal.pone.0066134.

Tobudic, S. et al. (2023) ‘The accelerated waning of immunity and reduced effect of booster in patients treated with bDMARD and tsDMARD after SARS-CoV-2 mRNA vaccination’, Frontiers in Medicine, 10, p. 1049157. 10.3389/fmed.2023.1049157.

Vazquez, M.I., Catalan-Dibene, J. and Zlotnik, A. (2015) ‘B cells responses and cytokine production are regulated by their immune microenvironment’, Cytokine, 74(2), pp. 318–326. 10.1016/j.cyto.2015.02.007.

Voysey, M. et al. (2021) ‘Safety and efficacy of the ChAdOx1 nCoV-19 vaccine (AZD1222) against SARS-CoV-2: an interim analysis of four randomised controlled trials in Brazil, South Africa, and the UK’, The Lancet, 397(10269), pp. 99–111. 10.1016/S0140-6736(20)32661-1.

Wagar, L.E. et al. (2021) ‘Modeling human adaptive immune responses with tonsil organoids’, Nature Medicine, 27(1), pp. 125–135. 10.1038/s41591-020-01145-0.

Yin, Q. et al. (2023) ‘A TLR7-nanoparticle adjuvant promotes a broad immune response against heterologous strains of influenza and SARS-CoV-2’, Nature Materials, 22(3), pp. 380–390. 10.1038/s41563-022-01464-2.

Zhang, H. et al. (2017) ‘A distinct subset of plasmacytoid dendritic cells induces activation and differentiation of B and T lymphocytes’, Proceedings of the National Academy of Sciences, 114(8), pp. 1988–1993. 10.1073/pnas.1610630114.

Zhong, C. et al. (2021) ‘Type I interferon promotes humoral immunity in viral vector vaccination’, Journal of Virology. Edited by G. Silvestri, 95(22), pp. e00925–21. 10.1128/JVI.00925-21.

Zhu, J., Huang, X. and Yang, Y. (2007) ‘Type I IFN signaling on both B and CD4 T cells is required for protective antibody response to adenovirus’, The Journal of Immunology, 178(6), pp. 3505–3510. 10.4049/jimmunol.178.6.3505.

